# Chorismate and isochorismate turnover reveal hidden dynamics of aromatic metabolism in Arabidopsis

**DOI:** 10.64898/2026.05.09.724013

**Authors:** Scott A. Ford, Anya E. Franks, Anya Hu, Christina De Melo, Sonia E. Evans, Matthew E. Bergman, Michael A. Phillips

## Abstract

The shikimate pathway provides precursors for phenolic metabolites in plant primary and secondary metabolism. Carbon flux measurements in intact Arabidopsis leaves using a novel isotopologue MS/MS methodology revealed unexpected dynamics between chorismate and isochorismate pools. ^13^CO_2_ labeling kinetics point to chloroplast-derived shikimate as a major bifurcation point, with approximately one third diverted away from the chloroplast. In contrast, the small pool of chorismate was almost exclusively chloroplast-localized and turned over rapidly, reaching more than 70% labeling within minutes. Isochorismate, a precursor to salicylic acid (SA) in the Brassicales, was present at 50-fold molar excess over chorismate but labeled much more slowly and appeared primarily extra-chloroplastic. Total isochorismate declined by 90% in the Arabidopsis *enhanced disease susceptibility* mutant, which lacks the isochorismate exporter. Non-aqueous fractionation further supported a primarily chloroplast-localized chorismate pool but extra-plastidic isochorismate. *Populus trichocarpa* and *Nicotiana benthamiana* leaves contained chorismate but only trace isochorismate, consistent with their use of the benzoyl-CoA route to SA. Carbon commitment calculations indicated that one third of the total chorismate pool in Arabidopsis leaves is diverted to isochorismate. Global leaf calculations based on elemental analysis-isotope ratio mass spectrometry and targeted isotope recovery indicate that only 0.03% of total assimilated carbon (5.27 pmol·mg^-1^ D.W.·min^-1^) enters the shikimate pathway in photosynthetically active mesophyll. For comparison ∼0.05% (7.75 pmol·mg^-1^ D.W.·min^-1^) enters the 2-*C*-methyl-D-erythritol 4-phosphate pathway, which provides precursors for photosynthetic pigments and electron carriers. The compartmentalization and turnover dynamics of shikimate, chorismate, and isochorismate suggest continuous demand for aromatic precursors in mesophyll tissue is comparable to demand for MEP-pathway derived pigments.

## Introduction

Chorismate is a central metabolic intermediate in plants, fungi, algae and bacteria, linking primary metabolism to an extensive array of aromatic products (El-Azaz and Maeda, 2025). As the product of the shikimate pathway (Maeda and Dudareva, 2012), chorismate represents the last common precursor for aromatic amino acids (AAAs; Phe, Tyr, and Trp) and numerous downstream metabolites, including phenylpropanoids, lignin, salicylic acid (Rekhter et al., 2019), tetrahydrofolate, phylloquinone, and naphthoquinones (Figure 1A). It is synthesized from D-erythrose-4-phosphate (E4P) and two equivalents of phospho*enol*pyruvate (PEP) via sequential enzymatic steps of the shikimate pathway. In photosynthetic tissue, E4P derives from the Calvin–Benson-Bassham (CBB) cycle, while PEP is reimported from the cytosol by the PEP:phosphate translocator 1 (PPT1) (Streatfield et al., 1999) (Figure 1B). The pathway begins with the condensation of E4P and PEP by 3-deoxy-D-arabino-heptulosonate-7-phosphate (DAHP) synthase (Yokoyama et al., 2022) followed by a rearrangement to dehydroquinate, dehydration to dehydroshikimate, and reduction leading to shikimate. This in turn is followed by phosphorylation to shikimate-3-phosphate and condensation with a second unit of PEP to 5- *enol*pyruvyl shikimate-3-phosphate (EPSP), whereupon chorismate synthase then dephosphorylates EPSP to yield chorismate (Rekhter et al., 2019). The dominant pathway to Phe in the plastid involves isomerization of chorismate to prephenate by chorismate mutase (CM), transamination to arogenate by prephenate aminotransferase, and dehydration to Phe by arogenate dehydratase. Cytosolic Phe is also thought to arise in flowering plants through a cytosolic CM isoform to generate prephenate, which is followed by prephenate dehydratase- mediated conversion to phenylpyruvate and transamination to Phe using Tyr as amine donor (Yoo et al., 2013; Qian et al., 2019). However, direct evidence supporting the cytosolic formation of a chorismate pool (Lynch, 2022) has not been reported.

**Figure 1.**
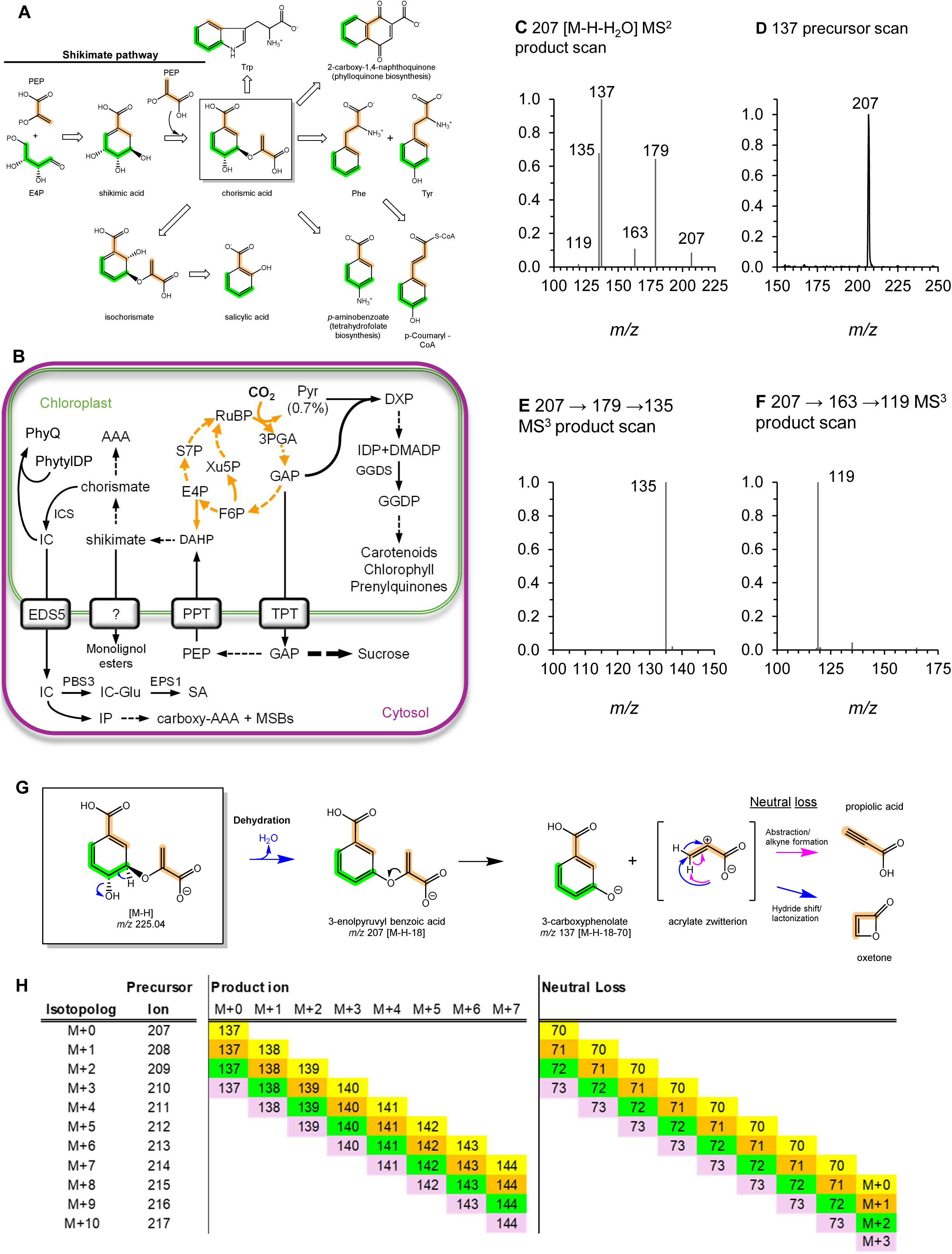
Roles of chorismate in primary metabolism and analysis of its ^13^C labeling pattern. **A**, Chorismate provides precursors for aromatic metabolites in primary plant metabolism. The contributions of PEP (orange) and D-erythrose-4- phosphate (E4P) (green) to the carbon skeletons of downstream phenolic metabolites are highlighted. Hollow arrows signify multiple enzymatic steps. OP signifies a phosphate moiety. **B**, Schematic highlighting the trafficking and metabolism of phenolics and terpenoids in Arabidopsis mesophyll described in this study. Dashed lines indicate multiple steps. Abbreviations: RuBP, ribulose-1,5-bisphosphate; 3PGA, 3-phosphoglycerate; GAP, glyceraldehyde-3- phosphate; F6P, fructose-6-phosphate; S7P, sedoheptulose-7-phosphate; xylulose-5-phosphate; Pyr, pyruvate; DXP, 1-deoxyxylulose-5-phosphate; IDP, isopentenyl diphosphate; DMADP, dimethylallyl diphosphate; PEP, phosphoenolpyruvate; E4P, erythrose-4-phosphate; PhyQ, phylloquinone; PhytylDP, phytyl diphosphate; TPT, triose phosphate/phosphate translocator; PPT, phosphoenolpyruvate/phosphate translocator; GGDS, Geranylgeranyl diphosphate synthase; ICS, isochorismate synthase; IC, isochorismate; EDS5, isochorismate translocator; PBS3, AvrPphB Susceptible 3; IC-Glu, isochorismate-9-glutamate; EPS1, enhanced pseudomonas susceptibility; SA, salicylic acid; IP, isoprephenate, cAAA, carboxy aromatic amino acids; MSB, *meta*-substituted benzoates. **C-H,** MS^2^ and MS^3^ spectra of chorismate and strategy for calculating fractional labeling in whole plant ^13^CO_2_ time course experiments. All scans were performed using direct infusion of an authentic chorismate standard at 10 µg·mL^-1^ (see Methods for details). The quasimolecular [M-H]^-^ ion (*m/z* 225) is unstable and undergoes in-source dehydration. **C**, Product scan of the in-source dehydration product of chorismate (*m/z* 207) showing the major products (*m/*z 137, 179, 135, 163, 119). **D**, The 207→179 MS^3^ scan, which implies loss of 28 Da from the 207 ion, is more likely the result of CO_2_ loss, O_2_ adduct formation, and loss of O. **D**, The 137 precursor scan confirmed that 207 is the exclusive precursor. **F,** MS^3^ scans verified that the *m/z* 163 and 119 product ions are the result of successive CO2 losses from the *m/z* 207 precursor ion. **G**, The principal MS/MS fragmentation of chorismate, involving loss of the C_3_ enolpyruvate group (*m/z* 207 → 137) from the insource dehydration product of chorismate, was selected to quantify ^13^C label in ^13^CO_2_ labeled plant extracts. **H**, Isotopologue expansion series used to track chorismate labeling during CID fragmentation. This multiplexed MRM table was required to quantify the relative abundances of the chorismate isotopolog precursor ions. Because of the distribution of carbon atoms in **G**, a total of 32 MRM transitions must be monitored simultaneously for exact calculation of isotopic labeling into chorismate and unambiguously account for all distributions of ^13^C atoms during CID.

While no shikimate transporter has been identified to date, a portion is thought to be exported from the plastid to the cytosol, where it participates as a re-usable esterification adaptor in monolignol biosynthesis. Hydroxycinnamoyl-CoA:shikimate/quinate hydroxycinnamoyltransferase esterifies *p*-coumaryl-CoA to *p*-coumaryl shikimate (Schoch et al., 2001; Hoffmann et al., 2003) which is hydroxylated to caffeoyl shikimate and subsequently re-esterified to CoA or hydrolyzed to caffeate (Vanholme et al., 2013). Phe exported from the plastid enters the phenylpropanoid pathway, initiated by phenylalanine ammonia lyase (PAL) (with contributions from tyrosine ammonia lyase in the Poaceae) (Barros and Dixon, 2020). Aromatic plant natural products derived from the shikimate pathway passing through these enzymatic steps reportedly account for up to 30% of carbon assimilated in woody species (Razal et al., 1996; Boerjan et al., 2003; Maeda and Dudareva, 2012).

Although chorismate is a precursor to multiple downstream pathways, its predominant plastidial role is supplying Phe through the arogenate pathway (Vogt, 2010). Much of this Phe is exported to the cytosol and converted to phenylpropanoids (Widhalm et al., 2015). In *Petunia hybrida*, chorismate conversion to Phe supports benzenoid and phenylpropene volatile biosynthesis (Qian et al., 2019). Cytosolic isoforms of chorismate mutase have been identified in several plant species (d’Amato et al., 1984; Singh et al., 1986; Westfall et al., 2014), suggesting chorismate may be present outside of the chloroplast. Several other enzymes of the shikimate pathway have duplicated paralogs in the cytosol in one or more plant species, leading to the ‘dual pathway hypothesis’ (Lynch, 2022), which posits that portions of the shikimate pathway are active in both compartments. Labeling studies indicate a significant extra-plastidic shikimate pool in Arabidopsis (Bergman et al., 2022), but analogous insights into chorismate trafficking remain lacking.

A major downstream fate of chorismate is isochorismate, particularly in Brassicales, where it supplies salicylic acid (SA), a central defense hormone. Chorismate is isomerized to isochorismate by isochorismate synthase (ICS) within the plastid, after which it exported to the cytosol by the transporter ENHANCED DISEASE SUSCEPTIBILITY 5 (EDS5). There it is conjugated to Glu by AvrPphB Susceptible 3 (PBS3) and subsequently hydrolyzed to SA (Torrens-Spence et al., 2019). The ICS-EDS5-PBS3 pathway evolved specifically within a lineage of the Brassicales order (Hong et al., 2025). In plants outside of this lineage, the bulk of SA is produced through the PAL pathway though cinnamate-derived intermediates (Liu et al., 2025; Ma et al., 2025; Wang et al., 2025; Zhu et al., 2025). This route may also contribute to SA formation in Arabidopsis (Huang et al., 2010). However, feeding Arabidopsis with ^13^C-labeled benzoic acid produced labeled SA whereas ^13^C-labeled Phe feeding did not (Wu et al., 2023). Thus, the contribution of the PAL pathway to SA production in Arabidopsis remains unresolved.

Despite this specialization of SA biosynthesis through the ICS pathway in the Brassicales, ICS homologs are conserved across land plants due to their essential role in phylloquinone biosynthesis in the chloroplast. Many species carry duplicated ICS paralogs, with varying functional divergence. In Arabidopsis ICS1 and ICS2 have overlapping functions, but ICS1 accounts for most SA production (Garcion et al., 2008). The *ics1/ics2* double mutant lacks phylloquinone but retains trace SA, consistent with limited contribution from the PAL-derived pathway (Wu et al., 2023). Outside of the Brassicales, ICS isoforms have diverged functionally. In cotton, ICS1 produces isochorismate for phylloquinone while ICS2 supplies SA (Guo et al., 2023), whereas in rice (Wang et al., 2024) and barley (Qin et al., 2019), isochorismate is required only for phylloquinone biosynthesis, while SA is produced through the PAL route.

The centrality of chorismate contrasts with how little is known about its turnover. Its instability, arising from a pre-aromatic ring structure and labile bonds, leads to rapid decomposition during MS analysis (Khera et al., 2017). Such fragmentation complicates quantitative assays of its metabolic role. Developing robust mass spectrometry methods would enable isotopic tracing, particularly with ^13^C, and thereby clarify flux through the shikimate pathway. Indeed, ^13^C labeling has become a standard tool in plant systems biology, improving models of primary and secondary metabolism (Ma et al., 2014; Wright et al., 2014; Xu et al., 2022).

^13^C labeling coupled to gas and liquid chromatography tandem mass spectrometry (GC-MS and LC-MS) has been instrumental in studying carbon flux in diverse metabolic networks (Szecowka et al., 2013; Ma et al., 2014; Xu et al., 2021; Bergman et al., 2022; Xu et al., 2024). Chorismate analysis by GC-MS is precluded by instability, unlike shikimate, whose isotopologues can be tracked with GC-MS using soft ionization (Bergman et al., 2022). Many central phosphorylated metabolites, such as intermediates of the 2-*C*-methyl-D-erythritol-4- phosphate pathway (MEP), are more tractable in MS/MS because their fragmentation yields dominant phosphate (*m/z* 97) and phosphite (*m/z* 79) ions devoid of carbon, simplifying isotopologue calculations (Heise et al., 2014; Wright et al., 2014; Evans et al., 2022). Their isotopologue distribution can thus be quantified using multiple reaction monitoring (MRM) simply by varying precursor mass (Wright and Phillips, 2014). In contrast, non-phosphorylated metabolites, including amino acids (Cocuron et al., 2017) and organic acids (Koubaa et al., 2014), often split carbon fragments during collision induced dissociation (CID), complicating isotopic analysis. Chorismate, with its multiple labile bonds, exhibits diverse CID fragmentation (Khera et al., 2017). Systematic characterization of its MS/MS behavior would allow isotopologue deconvolution in ^13^C labeling experiments. We therefore analyzed its fragmentation by tandem MS to support flux studies into its synthesis and transport.

In this report, we characterize the turn-over and transport of chorismate and isochorismate in Arabidopsis by implementing a multiplexed LC-MS/MS multiple reaction monitoring (MRM) technique. We determined that a substantial portion of chorismate is diverted to isochorismate, which is exported from the chloroplast and remains largely metabolically inert in the absence of biotic stress. We further report that the allocation of carbon to the shikimate pathway is a small fraction of total assimilated carbon in photosynthetically active mesophyll tissue that is comparable in magnitude to carbon delivered to the MEP pathway for production of terpenoid pigments. Both precursor pathways combined consume less than 1% of total carbon, underscoring the metabolic specialization of photosynthetic cell types and the contrasting allocation of carbon resources needed to meet developmental and physiological demands.

## Materials and Methods

### Plant cultivation and isotopic labeling

*Arabidopsis thaliana* (ecotype Columbia 0) wild-type or *mutant* seeds were introduced to moist BX soil (Promix) in 5 cm pots and incubated for three days at 4 °C in the dark, before being transferred to short day (SD) conditions (8/16 hour light/dark cycles at 150 µmol photons·m^-2^·s^-1^, 21 °C, and 60-70% relative humidity). Plants were watered as needed and fertilized once weekly with MiracleGro™ (20/20/20) according to manufacturer’s instructions.

Rosette stage Arabidopsis plants 6-8 weeks old were subjected to gas exchange measurements and ^13^CO_2_ *in vivo* labeling. This was accomplished with a hybrid LI-COR 6800 photosynthesis system fitted with a custom cuvette adaptor and an environmentally controlled dynamic flow chamber (Evans et al., 2022). Photosynthetically active surface area of plants was determined in Photoshop by comparison to size standards. Air flow containing CO_2_ at 400 µL·L^-1^ was passed over the enclosed rosette at 680 µmol air ·s^-1^ (∼1 L·min^-1^ at 21 °C) using the LI-COR console until the assimilation rate had stabilized, indicating a photosynthetic steady state (∼30 min). Labeling was initiated by rapidly switching from air supplied by the LI-COR 6800 to a pressurized gas tank (Linde Canada) containing synthetic air (80% N_2_, 20% O_2_) supplemented with ^13^CO_2_ at 400 µL·L^-1^. Relative humidity of the synthetic air was ensured by passage through a wash bottle prior to reaching the plant, and the exposure time of each time-course plant was monitored with a stopwatch until quenching by liquid nitrogen. The labeling time was corrected to the half-way point of the two atmospheres, as judged by the decay of the CO_2_ gas analyzer signal as ^13^CO_2_-containing air entered the chamber, usually in the range of 5-10 s. A total of 38 wild-type plants were labeled from times ranging from 5 s to 155 min.

Arabidopsis mutant lines were obtained from the Arabidopsis Biological Resource Centre (https://abrc.osu.edu/). This includes mutant lines of ISOCHORISMATE SYNTHASE 2 (*ics2*) (SALK_084635; AT1G18870), ISOCHORISMATE SYNTHASE 1 (*sid2-1*; AT1G74710), (Wildermuth et al., 2001), and ENHANCED DISEASE SUSPECTIBILITY 5 (*eds5-1*; AT4G39030). Genotyping primer sequences are listed in Supplemental Table 4.

*Nicotiana benthamiana* were grown under long day conditions (16/8 hour light/dark cycles) in environmentally controlled chambers with 300 µmol photons·m^-2^·s^-1^, 25 °C, and 60- 70% relative humidity. *Populus trichocarpa* leaves were harvested on the University of Toronto - Mississauga campus at midday in July 2025 and immediately flash frozen in liquid nitrogen for processing.

### Plant processing and extraction

Flash frozen, labeled plant tissue (or unlabeled control) derived from a single labeling experiment was ground to a fine powder in liquid nitrogen and lyophilized to dryness against a vacuum of 25 µbar for 24 h. The resulting powder was stored in a 2 mL tube at -20 °C until extracted. All labeled plant samples were analyzed by elemental analysis - isotope ratio - mass spectrometry (EA-IRMS) analysis as previously described to obtain the ^13^C/^12^C ratio and carbon content of bulk tissue (Bergman et al., 2021).

To extract polar metabolites for LC-MS/MS analysis, the sample was kept at 4 °C for all steps of the procedure. A 15 mg aliquot of dry tissue was weighed out on an analytical balance, extracted twice with 50% acetonitrile containing 10 mM ammonium acetate pH 9.0, and centrifuged at 16,000 g for 10 min after each extraction. An internal standard (200 ng deoxyglucose-6-phosphate) (Millipore-Sigma) was added at the beginning of the protocol. The extracts were lyophilized overnight, resuspended in 100 µL aqueous 10 mM ammonium acetate pH 9.0, and back extracted against 1 vol CHCl_3_. Following centrifugation at 16,000 *g*, the upper aqueous phase was diluted with 1 vol acetonitrile, filtered through a 0.2 µm PVDF InnoSep™ Spin filter (Canadian Life Science), and transferred to a 2 mL glass LC vial with a polypropylene insert.

For quantitation of MEP and shikimate pathway intermediates, standard curves were prepared using the protocol above with differing amounts of authentic standards in solvent only while maintaining internal standards constant (Supplemental Figure 5). To infer matrix effects, standard curves were then repeated in the presence of 15 mg control tissue with standards added at the beginning of the protocol using the same levels. For comparison, a calibration series was prepared in parallel where standards were added just prior to injection (Supplemental Figure 6). The results were compared to calculate recovery, matrix effects, and ion suppression using previously reported methods (Nasiri et al., 2021; Raposo and Barceló, 2021; Williams et al., 2023).

Sucrose analysis was carried out on 2 mg lyophilized tissue extracted with 500 µL 75% methanol (*v/v*) containing the same internal standards, centrifuged and filtered as above. Its concentration and isotopologue profile were analyzed by LC-MS/MS as described in Supplemental Table 2.

For gas chromatography-mass spectrometry (GC-MS) analysis of isotopically labeled plant tissue, 75% (*v/v*) methanolic extracts of a 5 mg tissue aliquot were dried on a nitrogen evaporator, resuspended in 100 µL pyridine, and converted to their methyloxime/trimethylsilyl (MeOX/TMS(derivatives in a two-step sequence as described previously (González-Cabanelas et al., 2015). To calculate label incorporation into shikimate by GC-MS, MeOX/TMS derivatized extracts were analyzed by ammonia positive chemical ionization as described previously (Bergman et al., 2022).

### Liquid chromatography - mass spectrometry analysis

LC-MS/MS targeted metabolite analysis reported was performed on an Agilent 1290 series II ultrahigh pressure LC, coupled to a Sciex 4500QTrap linear ion trap-triple quadrupole mass spectrometer. All metabolites described in this study were separated by hydrophilic interaction chromatography (HILIC) using an XBridge Premier BEH amide column (2.5 µm particle size, 2.1 mm × 150 mm; Waters Corporation) based on Gonzalez-Cabanelas et al. (González-Cabanelas et al., 2016) with minor modifications. The LC mobile phases consisted of 20 mM ammonium bicarbonate, adjusted to pH 10.5 with NH_4_OH (A) and 80% acetonitrile with 20 mM ammonium bicarbonate pH 10.5 (B). The flow rate was constant at 0.5 mL·min^-1^. The solvent gradient was as follows: 0-16% A (0-5 min), 16%-40% A (5-10 min), hold 40% A (10-15 min), and then re-equilibrate at 0% A (15-30 min).

^13^C labeled isotopologues of intermediates from the MEP and shikimate pathway detected in plant extracts were quantified by multiple reaction monitoring (MRM) using electrospray ionization in negative mode. Detailed data acquisition parameters, including mass transitions of isotopologue series for all analytes in this study, are listed in Supplemental Tables 2 and 5. The detection of analytes was optimized by compound optimization with purified standards to establish optimal ionization and fragmentation energies, and product and precursor scans were performed to select the best Q_1_/Q_3_ pairs for overall sensitivity and linearity.

Collision-induced dissociation (CID) of chorismate was investigated by direct infusion of a purified standard (Millipore Sigma, ≥98% purity) dissolved in a mixture of solvents A and B. The analyte was ionized in negative electrospray ionization (ESI) mode (-4500 V, 200 °C temperature). Precursor and product ion scans were acquired for the deprotonated parent ion ([M–H]⁻, *m/z* 225) and for the dehydrated in-source fragmentation product (*m/z* 207). No significant adduct ions were detected. To confirm structural identity of product ions, fragmentation pathways were further characterized with MS^2^ and MS^3^ using the linear ion trap function of this instrument, focusing on major product ions (*m/z* 179, 163, 153, 137, 135). Ion trap experiments using purified chorismate were performed using -16 V collision energy, dynamic fill time, and a mass range of *m/z* 50 – 200. Spectra were averaged over three replicate scans to improve the signal-to-noise ratio. All experiments were performed in triplicate to ensure reproducibility. These experiments defined the principal fragmentation routes of chorismate under negative CID, providing mechanistic insight into its MS behavior needed for its targeted detection in complex biological extracts.

### Calculation of fractional ^13^C enrichment and isotope recovery

Fractional labeling (F) of PEP and MEcDP were carried out as previously reported (Evans et al., 2024). Fractional labeling of chorismate and shikimate was determined using a newly developed multiplex MRM approach that accounts for all possible isotopologues and their distributions into product and neutral loss fragments during CID. When carbon atoms are retained in both the product ion *p* and neutral loss *l* following CID, the total transitions that must be monitored for an exact calculation of fractional labeling (F) must take into consideration all possible product ions that can arise from a given isotopolog precursor, based on physically possible distributions of ^13^C atoms between the product ion and neutral loss fragment. Because both the fragment ion and the neutral loss retain subsets of the total carbon skeleton in the case of chorismate, each isotopologue of the precursor (M+0 to M+10) can produce multiple product ions with distinct isotopic compositions. For chorismate, the *m/z* 207 → 137 transition was selected to analyze label incorporation in Arabidopsis leaves. All 10 carbons of the original chorismate molecule are present in the *m/*z 207 ion, while the product contains 7. A total of 32 unique transitions were required to account for all possible distributions of ^13^C in the 11 chorismate isotopologues (Figure 1H). By summing the integrated peak areas of the product ion isotopologues derived from each precursor isotopologue, we calculated relative abundances of each isotopologue. Following normalization to the monoisotopic peak, calculation of fractional ^13^C enrichment in the chorismate pool of ^13^CO_2_ labeled plants was accomplished using the following relationship:

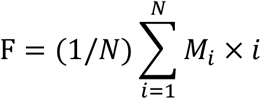

where *N* is the number of carbons of the compound of interest, and *M*_i_ is the fractional abundance of the *i*th isotopologue.

To calculate global carbon commitment and pathway specific carbon allocation, we first computed the carbon commitment (CC) for each individual plant at each timepoint into each metabolite, defined as the nmol of ^13^C atoms in each metabolite pool and calculated as follows:

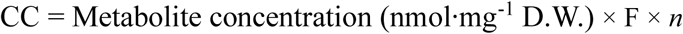

F represents the fractional labeling of that individual sample and *n* is the number of carbons in the metabolite. We next established the linear regression for carbon assimilation into each metabolite pool surveyed here using time-course data. The initial slopes of hyperbolic labeling curves for chorismate, PEP, and MEcDP were inspected for linearity, and time points ranging from 0 – 8 minutes presented R^2^ values of 0.97, 0.88, and 0.91, respectively. By plotting CC against time for the linear portion, we obtained their initial slopes. Once all slopes had been converted to the common units of pmol ^13^C atoms·mg^-1^ D.W. ·min^-1^, we obtained percent-wise commitment of assimilated carbon to each metabolite pool by dividing each slope by the total ^13^C assimilation slope derived from EA-IRMS analysis of all time-course labeled plants. The resulting dimensionless fraction represents the proportion of all assimilated carbon entering that metabolite pool during time-course labeling experiments.

### Enzymatic production of IC

To prepare an IC standard for LC-MS/MS analysis, the Arabidopsis ICS1 gene (AT1G74710) was PCR amplified without a chloroplast targeting peptide using Gateway attB primers (Supplemental Figure 3 and Supplemental Table 4). The PCR amplicon was cloned into the pDONR207 entry vector with BP Clonase II (Invitrogen^TM^) and then transferred to a Gateway^TM^ modified pET28 vector (Yu and Liu, 2006) for bacterial protein expression fused to a His_9_ affinity tag using LR Clonase II. Chemically competent BL21 (DE3) *Escherichia coli* cells were transformed with the expression vector containing the truncated ICS construct and plated on Luria-Bertani agar media supplemented with kanamycin (50 μg·mL^-1^) for overnight incubation at 37 °C. Positive colonies were selected by colony PCR, grown in 5 mL starter cultures overnight at 37 °C, 220 RPM, and each used to inoculate a separate 200 mL culture of Terrific Broth with the selective antibiotic, kanamycin (50 μg·mL^-1^). Cultures were incubated under previous conditions until an optical density at 600 nm (OD_600_) of 0.6-0.8 was achieved, after which cultures were induced for protein expression with 1 mM isopropyl β-D-1- thiogalactopyranoside (IPTG), and the temperature reduced to 16 °C for an incubation period of 18-20 hours at 230 RPM. Cells were harvested by centrifugation at 4 °C and 1,000 *g* for 30 minutes, after which the cell pellets were resuspended in 10 mL of lysis buffer (phosphate buffered saline, 60 mM imidazole, 1 mg·mL^-1^ lysozyme) with a protease inhibitor cocktail mix (Sigma-Aldrich®) used according to the manufacturer’s instructions. The resuspension was incubated on ice for 30 minutes, subjected to two freeze-thaw cycles, and sonicated at 60% power for 6 cycles of 10 seconds power and 10 seconds of an intermittent cooling period. The suspension was clarified by centrifugation for an hour at 4 °C, 4,000 *g*, and the supernatant purified by Ni-NTA agarose (QIAGEN) chromatography as per the manufacturer’s instructions. Eluent was desalted with desalting buffer (50 mM Tris-HCl pH 8.0, 10 mM MgCl_2_, 150 mM NaCl, 10% glycerol) using an Amicon® ultra centrifugal filter (10 kDa MWCO; Millipore), and the protein concentration estimated using a standard Bradford assay.

Assays were performed in duplicate 100 μL reactions with 0.8 µg of ICS1 per assay in desalting buffer supplemented with 10 μM chorismate and incubated at 25°C for an hour. Reactions were terminated by vortexing with one vol of CHCl_3_, centrifuged for 10 minutes at 10,000 *g*; after which 50 μL of the aqueous supernatant was diluted with 50 μL of acetonitrile and transferred to a 2 mL LC vial with a polypropylene insert for analysis by LC-MS/MS.

### Data analysis

Sciex OS (v2.0) and Agilent MassHunter Qualitative Analysis (v10.0) were used to analyze LC-MS/MS and GC-MS data, respectively. Analyte peaks were identified by matching retention times and masses to authentic standards, then peak areas were determined by integrating extracted ion chromatograms (EIC). Absolute quantification of metabolites was determined by normalizing peak areas to internal standards and comparison to an external calibration curve of authentic standards. Recovery of internal standard was calculated by comparing DGP peak areas in non-tissue controls to DGP recovered in Arabidopsis leaf extracts (15 mg wild-type Arabidopsis tissue) prepared under the same conditions. To assess statistical significance, a two-tailed student’s *t*-test was used with a significance threshold set to p < 0.01. Time course-labelling curves were generated using ^13^C mass spectral data as described elsewhere (Evans et al., 2024).

### Non-Aqueous Fractionation

Subcellular metabolite distribution was analyzed by non-aqueous fractionation using the benchtop protocol of Fürtauer et al. (Fürtauer et al., 2016). Approximately 25 mg of finely ground lyophilized tissue was resuspended in 2 mL chilled heptane:tetrachloroethylene (TCE) (0.484:1, v/v; ρ = 1.3 g/cm³) and further homogenized on ice using a Tenbroek homogenizer. The homogenate was transferred to a 50 mL tube, and the homogenizer was rinsed with 8 mL of the same solvent, which was pooled with the sample.

The suspension was sonicated in an ice-water bath for 10 min, filtered through a 20 μm nylon mesh, and the filter washed with 10 mL cold heptane. The filtrate was centrifuged (20 min, 4000 *g*, 4 °C), the supernatant discarded, and the pellet resuspended in 1 mL TCE and transferred to a 2 mL tube. Following an additional 10 min cold sonication, samples were centrifuged (12.5 min, 16,000 *g*) and the pellet retained to yield the densest fraction. The supernatant was sequentially diluted with heptane to decrease density, followed by sonication (10 min) and centrifugation (12.5 min, 16,000 *g*) at each step to generate pellets of decreasing density relative to the initial fraction. This iterative process produced fractions corresponding to densities of approximately 1.33, 1.41, 1.49, 1.61 g/cm^3^ (F2-F5; Supplemental Figure 4).

Each pellet was resuspended in 900 μL heptane:TCE adjusted to its respective density, split into two aliquots, washed with 1 mL heptane, and centrifuged (12.5 min, 16,000 *g*). Pellets were dried under nitrogen and stored at −20 °C. The final supernatant fraction was similarly split, dried, and stored. After collecting the lowest-density pellet (F2; ρ = 1.33 g/cm^3^), the remaining supernatant, containing material less dense than 1.33 g/cm^3^ (“F1”) was split into two equal aliquots, dried under nitrogen, and stored at -20 °C prior to metabolite analysis. For polar metabolite analysis, dried fractions were resuspended in 100 μL cold 10 mM ammonium acetate (pH 9.5), extracted with an equal volume of chilled chloroform, vortexed (20 min, 4 °C), and centrifuged for phase separation. A 50 μL aliquot of the aqueous phase was mixed with 50 μL cold acetonitrile and analyzed by HILIC LC–MS/MS as described above. For organic acids, samples were diluted 1:100 in acetonitrile:10 mM ammonium acetate (1:1, v/v; pH 9).

Relative metabolite distributions (*n* = 3 biological replicates) were calculated as the percentage of total peak area across the five fractions. Statistical analyses of metabolite data were performed using MetaboAnalyst 6.0 and Python scripts utilizing the scikit-learn (v1.6.1), and scipy (v1.16.3) libraries. PCA (autoscaled data) and hierarchical cluster analysis (HCA; Euclidean distance, average linkage) were used to assess fraction resolution and reproducibility (Beshir et al., 2019). HCA dendrograms were generated using Seaborn (v0.13.2) and matplotlib (v3.10.0) in Python. Clustering performance was quantified using the Normalized Mutual Information (NMI) score calculated for a fixed 5-cluster model (corresponding to the five fractions) using the `metrics.normalized_mutual_info_score` function in scikit-learn.

Pairwise Pearson correlation coefficients (r) were calculated for metabolite distributions across the five density fractions (1.61, 1.49, 1.41, 1.33, and <1.33 g/cm³), averaged across replicates, and visualized as a symmetric hierarchical clustermap (Euclidean distance, single linkage). Correlation strength and clustering pattern were used to infer subcellular co- localization. Compartmental distributions (chloroplast, cytosol, vacuole) were estimated using NAFalyzer (Hernandez et al., 2023). DXP and UDP-glucose were used as chloroplast and cytosolic markers, respectively, and the mean distribution of malate, fumarate, and citrate as a vacuolar marker (Szecowka et al., 2013). Outliers were handled as described by (Krueger et al., 2011). NAFalyzer output is provided in Supplemental Table 3.

## Results

### Resolving the cryptic fragmentation of chorismate enables flux measurements of the shikimate pathway

We surveyed the labeling and turnover of shikimate and chorismate by ^13^C-LC-MS/MS to support flux and transport studies of the shikimate pathway in plant cells. Their fragmentation patterns under CID were characterized to determine the best precursor/product pairs for monitoring label incorporation in ^13^CO_2_ treated plants. Shikimate undergoes a straightforward loss of CO_2_ and 2 water molecules during CID to form a resonance stabilized phenolate ion (*m/z* 173 → 93; Supplemental Figure 1), and this fragmentation pathway has been previously reported to quantify both pool size and label incorporation (Jaini et al., 2017; El-Azaz and Maeda, 2024).

Chorismate, in contrast, displays more complex behavior. Under negative ESI conditions, the chorismate M-H^-^ ion readily undergoes in-source dehydration to yield the aromatically stabilized 3-*enol*pyruvyl benzoate (*m/z* 225→207). During CID, the 207 precursor yields product ions of *m/z* 137, 179, 135, 163, and 119 in order of decreasing abundance (Figure 1C). Consistent with this, a precursor scan of the *m/*z 137 product ion confirmed that the *m/z* 207 ion is its exclusive source (Figure 1D), while the *m/*z 179 ion further fragmented to a product of *m/z* 135 (Figure E). Additional MS^2^ and MS^3^ scans indicate that two successive CO_2_ losses from 3- *enol*pyruvyl benzoate account for the *m/z* 207→163→119 fragmentation sequence (Figure 1F), ultimately forming a product whose identity is likely phenoxyvinylate ion (Supplemental Figure 1). The dominant transition (*m/z* 207→137) involves elimination of a 70 Da neutral fragment from 3-*enol*pyruvyl benzoate to 3-carboxyphenolate (Figure 1G). The neutral loss may be explained through elimination of an acrylate zwitterion, which can rearrange to propiolic acid or, less likely, smaller amounts of oxetone (Figure 1G).

The origin of the *m/z* 179 ion could, in principle, reflect concerted loss of CO_2_ and 2H· from the chorismate quasimolecular ion, [M-H-46]^-^ (Supplemental Figure 1E), leading to aromatization and retention of the phenol group. MS^3^ scans confirmed that the 225→179→153→109 sequence was readily detected, consistent with an initial loss of 46, corresponding to loss of CO_2_ and 2H·, followed by loss of acetylene and a second CO_2_, to produce 5-carboxyl-2-hydroxy phenolate (*m/z* 153) and 2-hydroxyl phenolate (*m/*z 109) (Supplemental Figure 1). However, precursor scans confirmed that 3-*enol*pyruvyl benzoate (*m/z* 207), and not deprotonated chorismate (*m/z* 225), was the main precursor for the *m/z* 179 product ion (Supplemental Figure 2A-C). A neutral loss of 28 implied by the 207 → 179 mass transition suggests loss of either carbon monoxide or ethylene, neither of which could be readily explained considering the likely neutral leaving groups of 3-*enol*pyruvyl benzoate. Based on similar observations in benzoic acid (Liu et al., 2024), we considered O_2_ adduct formation due to trace contaminants followed by loss of neutral O from the ensuing peroxide for a net loss of 28 Da (Supplemental Figure 1D). This was confirmed with MS^3^ ion trapping scans, which suggested that the *m/*z 179 ion may further lose CO_2_ (44 Da), followed by loss of ketene (42 Da) to form a phenolate ion (207→179→135→93; Supplemental Figure 2E). Based on the relative signal intensities and structural diagnostic value of the various chorismate fragmentation pathways, the *m/z* 207→137 MRM transition was selected for isotope incorporation calculations to characterize flux and localization of the shikimate pathway in Arabidopsis leaves.

### Chorismate and shikimate turnover in Arabidopsis leaves demonstrate pathway branching and transport kinetics

The dominant *m/z* 207→137 MS/MS fragmentation pathway of chorismate partitions carbon atoms between the neutral loss and product ions, complicating quantitative isotopologue analysis due to uncertainty in ^13^C distribution between fragments. To resolve this, we developed a multiplexed multiple-reaction monitoring (MRM) strategy enabling complete isotopologue coverage for chorismate and its fragment ions (Figure 1H). Using the full set of 32 MRM transitions derived from the monoisotopic 207→137 (C_10_→C_7_) transition, we quantified ^13^C incorporation into chorismate and related polar metabolites in Arabidopsis leaves exposed to ^13^CO_2_ under controlled physiological conditions for 4.8 s – 154.8 min prior to rapid freezing. The resulting time-resolved labeling curves exhibited hyperbolic kinetics for both PEP and chorismate, characterized by nearly identical apparent rate constants (*k* = 0.185 and 0.198 min^-1^, respectively), corresponding to half-lives of 5.41 and 5.06 mins and plateau enrichments of 68.4 % and 73.4 %, respectively, including natural abundance (Figure 2 and Supplemental Table 1). This similarity suggests a tight correlation between the labeled carbon entering chorismate and a rapidly equilibrating chloroplastic PEP pool.

**Figure 2.**
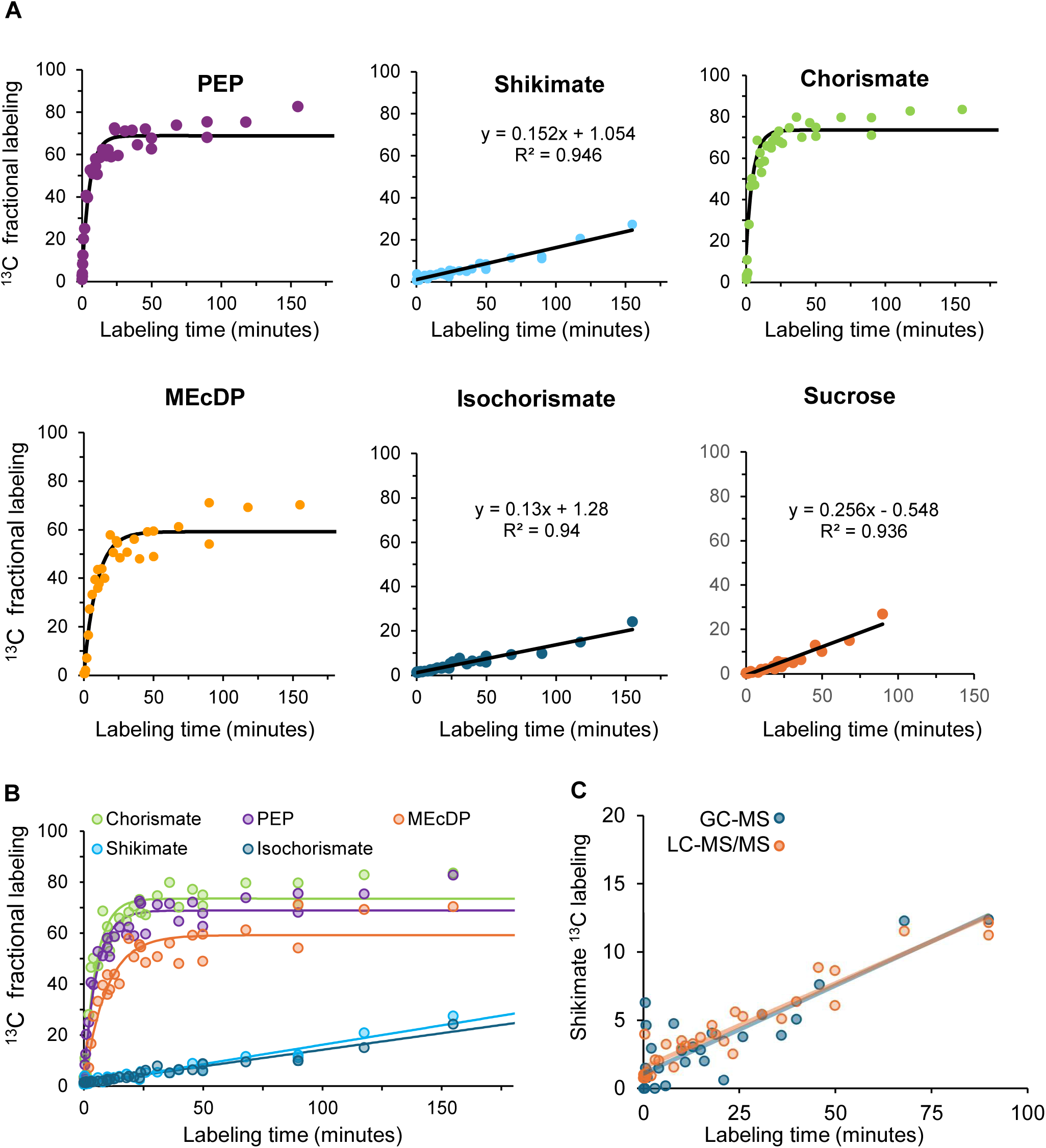
^13^CO_2_ time-course labeling of whole Arabidopsis rosettes under physiological conditions. **A**, Fractional labeling curves for select intermediates in the shikimate and methylerythritol phosphate pathways. For rapid (hyperbolic) labeling metabolites, curve fitting was performed using the exponential equation, *A* × (1 − e^[−k × *t*]^), where *A* is the labelling plateau, *k* is the kinetic rate constant, and *t* is the labelling time. For slow (approximately linear) labeling metabolites, a linear regression model was applied. For additional curve fitting parameters, see Table 1. **B,** Overlay of all fractional labeling time-course curves. **C**, GC-MS and LC-MS/MS techniques were compared for calculating shikimate labeling in intact Arabidopsis plants following ^13^CO_2_ labeling. Ammonia positive chemical ionization GC-MS quantification of methyloxime/ trimethylsilyl derivatized shikimate ^13^C isotopologues (Bergman et al., 2022) yielded % atom labeling results closely comparable to MRM LC-MS/MS measurements (See Methods). To calculate fractional labelling, we used the following equation: 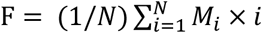; where *N* is the number of carbons of the compound of interest, and *M_1_* is the fractional abundance of the ith isotopologue.

**Table 1.** ^13^C incorporation rates into individual metabolite pools compared with total ^13^C in bulk leaf tissue following ^13^CO_2_ whole plant labeling. Initial slopes were determined from plants labeled for 0 – 7.82 mins, during which ^13^CO_2_ incorporation was approximately linear for all metabolites surveyed. Percent of total ^13^C was determined by dividing the initial slope of each metabolite by the slope of ^13^C incorporation into bulk leaf tissue, as determined by elemental analysis – isotope ratio mass spectrometry (EA-IRMS)^a^. Standard errors shown in parentheses.

| Pool | Pool size<br>(nmol · mg<sup>-1</sup>) | Initial slope<br>(pmol $^{13}\text{C}$ · mg<sup>-1</sup><br>D.W. · min<sup>-1</sup>) | % of Total $^{13}\text{C}$ | n |
| --- | --- | --- | --- | --- |
| Total $^{13}\text{C}$ (EA-IRMS) | n/a | 15162.21 (714.07) | 100 | 45 |
| MEcDP (MEP pathway) | 0.028 (0.0032) | 7.75 (0.43) | 0.051 | 13 |
| Shikimate (E4P + PEP) | 0.491 (0.031) | 5.27 (0.21) | 0.034 | 33 |
| Chorismate (E4P + 2PEP) | 0.005 (0.0004) | 4.74 (0.40) | 0.031 | 16 |
| Isochorismate (E4P + 2PEP) | 0.134 (0.010) | 1.67 (0.07) | 0.011 | 13 |
| Sucrose | 38.53 (4.13) | 1181.88 (23.99) | 7.795 | 28 |

<sup>a</sup> Bergman et al. (2021) *Plant Methods* 17 (1), 32

In contrast, shikimate displayed a low, linear rate of ^13^C incorporation under identical labeling conditions with a slope of 0.15% min^-1^ and a y-intercept of 1.05% (Figure 2). When the same samples were analyzed by soft ionization GC-MS, the molecular ion cluster of the 4× trimethylsilyl derivative of shikimate (*m/*z 480.2 – 487.2) gave nearly identical labeling results (Figure 2C). This was consistent with previously reported values obtained with this method (Bergman et al., 2022) and validated our LC-MS/MS MRM method for quantifying isotopologues. The intercept closely matched the expected natural ^13^C abundance of 1.07% for a C_3_ species, including ^13^C depletion (Bergman et al., 2021), validating both calibration and quantification procedures. The transitions used to calculate isotopic enrichments in shikimate and other analyzed metabolites are listed in Supplemental Table 2. Our analytical conditions were unable to quantify D-erythrose-4-phosphate in leaf extracts, and we were only able to infer its uptake into the shikimate pathway by comparing the labeling curves of PEP and shikimate.

Control metabolites confirmed the sensitivity of this approach to biochemical turnover rate. Sucrose, representative of slowly exchanging cytosolic carbon pools, exhibited a linear and gradual incorporation of ^13^C (Figure 2). Conversely, methyl-D-erythritol-2,4-cyclodiphosphate (MEcDP), an intermediate of the plastidic MEP pathway, showed rapid, pseudo first-order labeling kinetics consistent with its proximity to the primary site of ^13^CO_2_ assimilation (Evans et al., 2024).

To account for instability of intermediates during pool size calculations, we first conducted standard addition recovery assays by comparing peak intensities of authentic standards added to plant matrix either at the beginning of the extraction protocol or just before injection (Nasiri et al., 2021; Raposo and Barceló, 2021; Williams et al., 2023). These values were then used to calculate recoveries and correct metabolite concentrations derived from external calibration curves. Recoveries for chorismate, MEcDP, and shikimate were calculated at 82.4%, 71.4%, and 63.1%, respectively, while PEP recovery was <4%, consistent with previously reported values (El-Azaz and Maeda, 2024). The low recovery and instability of PEP precluded accurate assessment of its pool size by this approach, although this did not preclude determination of its labeling pattern.

Labeling and pool-size data were next combined to estimate absolute carbon fluxes into shikimate and chorismate as a function of time (Figure 3). Initial velocities for hyperbolically labeling metabolites were calculated from the linear portion of their exponential rise (0-8 min), while linear-labeling metabolites were analyzed directly by linear regression of their ^13^C incorporation slopes (Table 1). Comparing the initial velocity of carbon entering the shikimate pool (5.27pmol ^13^C atoms·mg^-1^ D.W.·min^-1^) to the shikimate-derived portion (7 of 10 carbon atoms) entering chorismate (0.7 × 4.74 = 3.32 pmol ^13^C atoms·mg^-1^ D.W.·min^-1^; Table 1), we estimate that 63% of the carbon entering shikimate continues to chorismate. If we assume shikimate is not converted to chorismate in the cytosol, this implies that ∼37% of all shikimate exits the shikimate pathway prematurely, remains metabolically inactive, and likely resides outside the chloroplast, a figure that comports well with its slow, linear rate of labeling.

**Figure 3.**
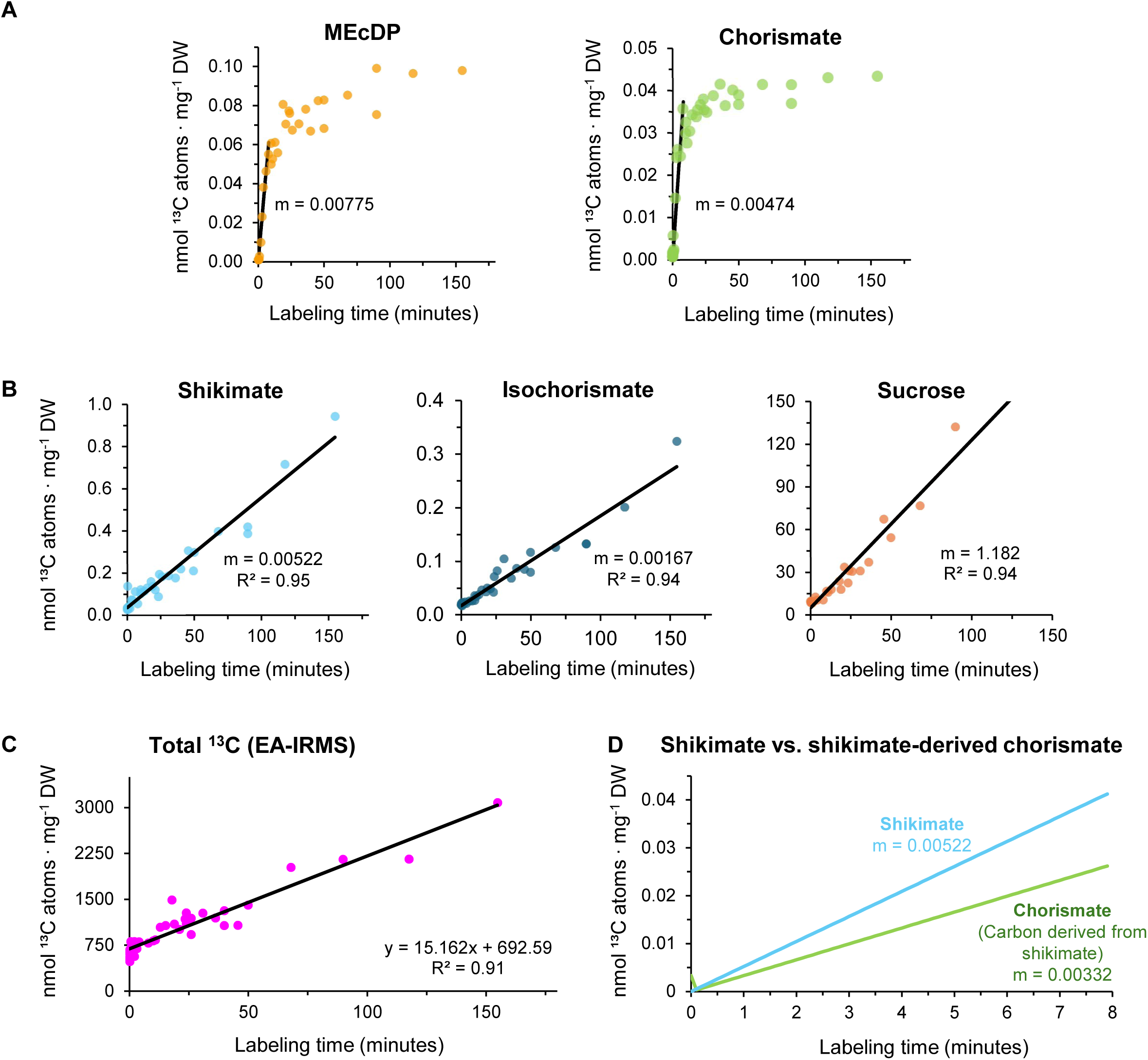
Carbon commitment to central metabolites compared to global carbon assimilation. Carbon commitment was calculated by multiplying the fractional labeling of each time point by the average pool size and the number of carbon atoms. Initial slope for metabolites displaying non-linear (**A**) or linear (**B**) labeling kinetics was obtained from the linear portions of the curves (0-8 minutes). Slopes of each metabolite were compared to the slope of total bulk tissue ^13^C enrichment, as determined by elemental analysis – isotope ratio mass spectrometry (**C**), to obtain a dimensionless fraction estimating the portion of total carbon directed to a specific metabolite pool. (**D**) A comparison of carbon commitment into shikimate and chorismate pools. Carbon commitment into chorismate was derived from the linear portion of the carbon commitment curve (0-8 minutes) and multiplied by the fraction of its carbon atoms that are derived directly from shikimate (7 of 10).

When the incorporation rates derived from LC–MS/MS were normalized to the total ^13^C assimilation rate measured by elemental analysis – isotope ratio mass spectrometry (EA–IRMS) (Figure 3C), the resulting ratios provided a measure of the fractional allocation of newly assimilated carbon through each metabolic pathway. This indicated that only ∼0.031% of total assimilated carbon completed the full shikimate pathway sequence to reach chorismate (Table 1), whereas 0.034% entered the shikimate pool, consistent with the inference that about a third of total shikimate enters a metabolically inactive or slowly recycled pool. Comparing this to the carbon allocation to an adjacent precursor pathway, analysis of the plastidial MEP pathway (using MEcDP as a representative intermediate) indicated that ∼0.05% of the total assimilated carbon was channeled into terpenoid precursor biosynthesis, or about 1.7 times the carbon observed entering the shikimate pathway. As a high-flux reference, the corresponding estimate for sucrose showed that during the middle portion of the day, about 8% percent of assimilated carbon (1,181.9 pmol ^13^C atoms·mg⁻¹ D.W.·min⁻¹) was diverted toward sucrose synthesis, consistent with its dominant role as the principal transport and storage carbohydrate (Figure 2 and 3; Table 1).

These results collectively imply that, in photosynthetically active mesophyll tissue, flux through the MEP pathway slightly exceeds that of the complete shikimate pathway but that the two precursor pathways are comparable in magnitude. This partitioning is physiologically plausible. The MEP pathway supports continuous synthesis and turnover of photosynthetic pigments derived from isopentenyl and dimethylallyl diphosphates on a timescale of hours (Beisel et al., 2010), whereas flux through the shikimate pathway must satisfy both protein production as well as vascular tissues (Ehlting et al., 2005) which prioritize Phe-derived lignification.

### A large, inactive isochorismate pool accumulates outside the chloroplast

Chromatographic separation of polar metabolites revealed that the small chorismate peak detected in Arabidopsis leaf extracts was preceded by a much more abundant isobaric compound (Figure 4A). It was approximately 50-fold larger than that of chorismate and exhibited similar MS/MS fragmentation behavior (*m/z* 207→137). Although prephenate is isobaric with chorismate, it cannot undergo spontaneous aromatization by in-source dehydration. CID of a prephenate standard confirmed a distinct transition (*m/z* 225→163) (Supplemental Figure 2F), corresponding to a concerted decarboxylation–dehydration event occurring after Q_1_ selection, consistent with observations by El-Azaz and Maeda (El-Azaz and Maeda, 2024). This diagnostic transition was absent in the *m/z* 225 product ion scan of chorismate (Supplemental Figure 2C), ruling out prephenate as the source of the major isobaric peak. We therefore hypothesized that the abundant isobar represented isochorismate, a positional isomer of chorismate and the immediate precursor of SA. Because a commercial standard was unavailable, isochorismate was synthesized *in vitro* using purified recombinant *Arabidopsis thaliana* ISOCHORISMATE SYNTHASE 1 (ICS1, At1g74710) expressed heterologously and assayed with chorismate as substrate (Supplemental Figure 3). The enzymatic product was analyzed by LC–MS/MS, and its retention time precisely matched that of the unknown isobaric peak in Arabidopsis leaf extracts, confirming the identity of the compound as isochorismate (Figure 4G). Further validation was obtained using Arabidopsis ICS knock-out lines. Chorismate pool size was significantly larger than wild-type in the *ics2* mutant (*p* < 0.005), consistent with the absence of the constitutively active ICS2. However, the decline in isochorismate levels was only statistically significant in the *sid2* mutant (*p* < 0.01). In *sid2-1* (*ics1*) and *ics2* single mutant backgrounds, isochorismate molar excess over chorismate compared to wild-type was reduced from 52.1 (±5.9) to 22.2 (±5.7) and 28.6 (±3.8), respectively (Figure 4B, C, and J). This change in isochorismate:chorismate ratio was statistically significant (*p* < 0.0005 and 0.005 for *ics2* and *sid2-1*, respectively, compared to wild-type) (Figure 4H-J). While the average isochorismate pool size decreased in each of the single knockout lines, it was not abolished entirely, consistent with the partial redundancy between the two ICS paralogs (Garcion et al., 2008).

**Figure 4.**
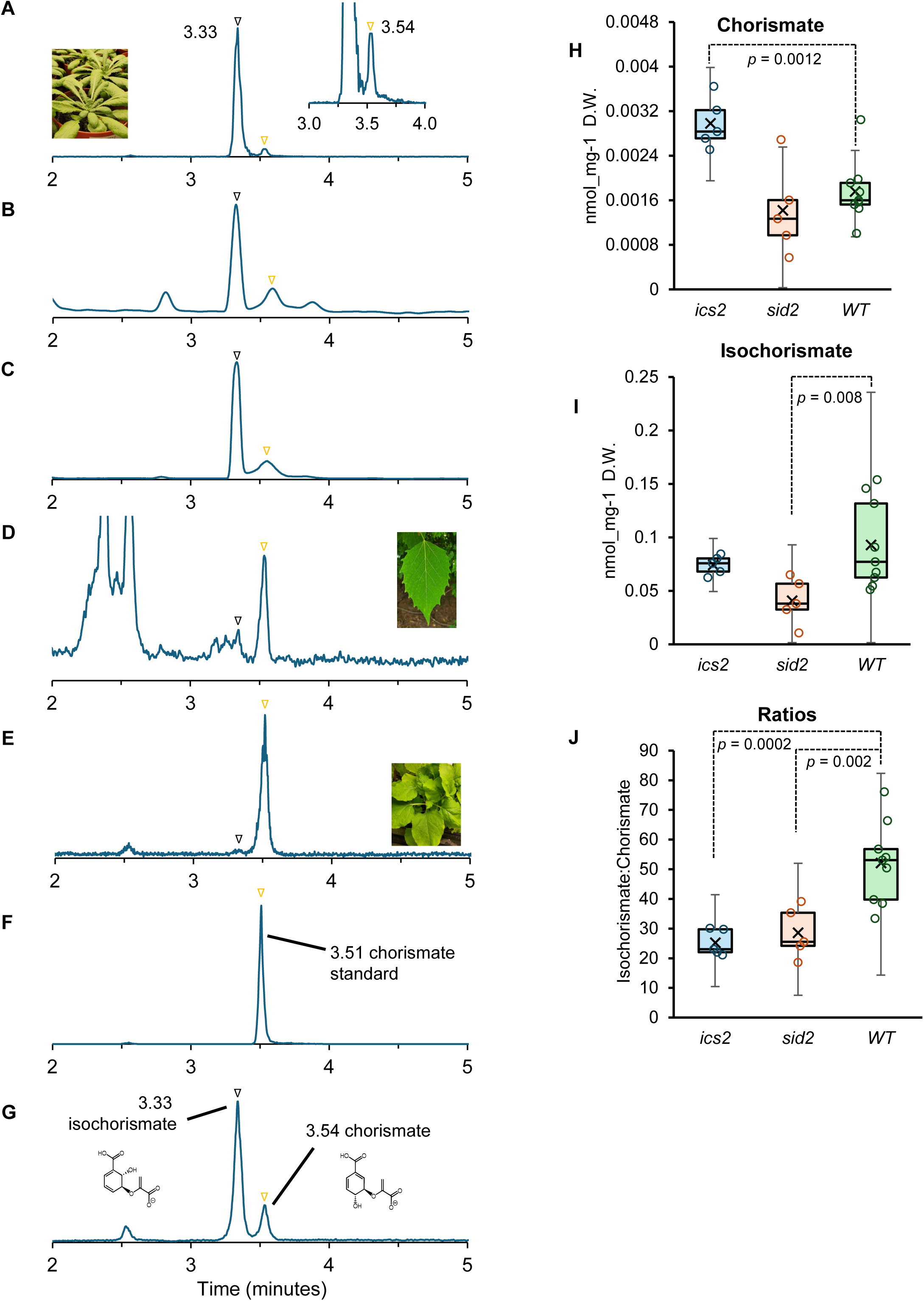
Detection of chorismate and isochorismate in whole leaf extracts by LCMS/MS. Polar metabolite leaf extract of **A**, Wild-type Arabidopsis; **B,** *sid2-1* mutant (SA INDUCTION- DEFICIENT2); **C,** *ics2* mutant (ISOCHORISMATE SYNTHASE 2); **D,** *Populus trichocarpa*; **E**, *Nicotiana benthamiana*; **F**, chorismate standard; **G**, enzymatically generated isochorismate standard (see Methods for additional details). The y axis represents the detector signal corresponding to the *m/z 2*07→137 mass transition (counts per second) from the presumed *m/*z 225 pseudomolecular ion, which undergoes an ion source fragmentation. Orange triangle = chorismate; black triangle = isochorismate; **H-I**, chorismate and isochorismate pool size in *ics2*, *sid2-1*, and wild-type (WT); **J**, pool size ratios of isochorismate:chorismate in ICS mutants and wild-type. P-values represents a two-tailed student’s t-test with n=5 for *ics2* and *sid2,* and n=9 for WT. Retention time drift between batches was corrected with an authentic chorismate standard.

We employed the same global isotope recovery and kinetic curve fitting approach to isochorismate, which exhibited a slow, linear labeling pattern resembling that of shikimate (Figure 2 and 3). This suggested that a large, metabolically stable isochorismate pool, likely outside the chloroplast, accumulates under basal conditions. We tested this idea with a time-course labeling series of the *enhanced disease susceptibility* 5 (*eds5*) mutant, which is defective in the isochorismate transporter (Nawrath et al., 2002). In the *eds5-1* mutant line, isochorismate pool size was only ∼10% of the wild-type plant (Figure 5A-B), consistent with a disruption of the export of isochorismate from chloroplasts. The linear labeling rate of this highly reduced isochorismate pool in the *eds5-1* mutant was nearly double that of the wild-type (Figure 5C), presumably reflecting the absence of the unlabeled isochorismate pool outside the chloroplast observed in wild-type plants.

**Figure 5.**
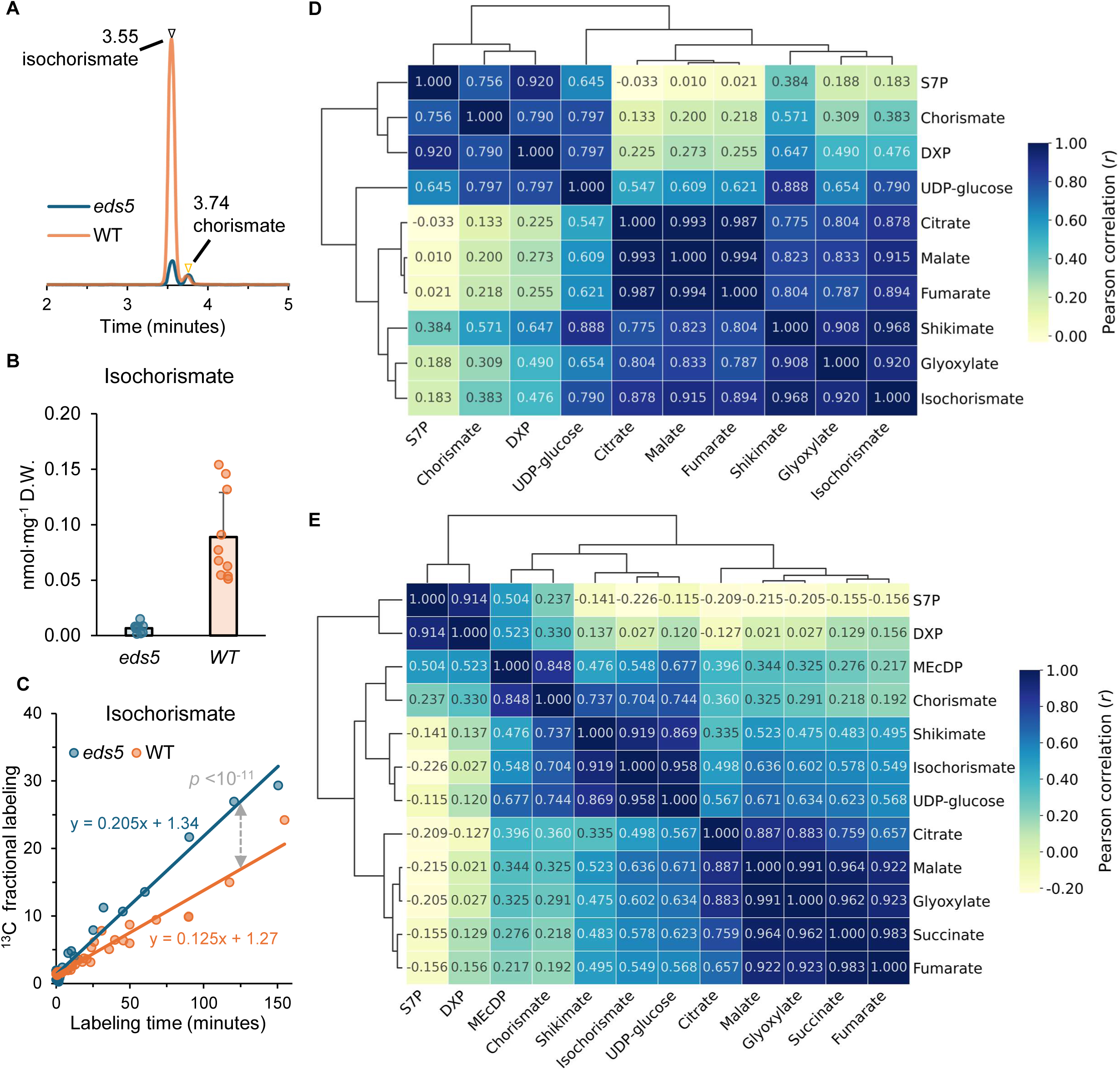
Metabolite profiling of the *eds5* mutant and subcellular distribution of selected metabolic intermediates. **A**, Chorismate (yellow triangle) and isochorismate (black triangle) detection in *eds5* and wild-type (WT) leaf extracts by LCMS/MS (y axis represents the detector signal corresponding to the *m/z 2*07→137 mass transition (counts per second)). **B,** Isochorismate steady-state concentration in illuminated Arabidopsis rosette tissue from wild-type and *eds5-1* mutant plants. The latter is defective in the isochorismate translocator. **C**, ^13^C time-course labeling of the isochorismate pool in intact wild-type and *eds5-1* individuals. The trend line differences are highly significant (p = 3.8 × 10^-12^) **D-E**, correlation analysis of metabolite distributions across five fractions of varying densities obtained through non-aqueous fractionation (see Methods and Materials for details). Consensus correlation matrix showing Pearson correlation coefficients *(r)* for pairwise metabolite combinations for **D** wild-type and **E** *eds5-1*. Following non-aqueous fractionation of Arabidopsis leaves, correlations were determined from metabolite distributions across the five obtained fractions with densities ranging from <1.33 to 1.62 g/cm³. The symmetric hierarchical clustered heatmap was generated using Euclidean distance and average linkage and depicts the averaged correlations of independent biological replicates (n=3). Abbreviations: S7P; sedoheptulose-7-phosphate; DXP, 1-deoxy-D-xylulose 5-phosphate; MEcDP, 2-C-methyl-D-erythritol-2,4-cyclodiphosphate; UDP-glucose; uridine diphosphate glucose.

To further investigate the compartmentation of chorismate and isochorismate, we conducted non-aqueous fractionation (NAF) using wild-type and *eds5-1* leaf tissue (Fürtauer et al., 2016). We targeted polar, diagnostic marker metabolites by hydrophilic interaction LCMS/MS to test the hypothesis that chorismate is largely or completely confined to the chloroplast while isochorismate is mostly stored outside that compartment following its export through EDS5. Using a heptane:tetrachloroethylene gradient, we analyzed plastidic, vacuolar, and cytosolic marker metabolites across density gradient fractions (Krueger et al., 2011; Szecowka et al., 2013) and calculated their Pearson correlation coefficients (Figure 5D-E). Quality control of the NAF data was performed using principal component and hierarchical clustering analysis (HCA) of metabolite distribution across fractions (Krueger et al., 2011) (Supplemental Figure 4). HCA of wild-type samples (*n* = 3) yielded a normalized mutual information (NMI) score of 0.81 for clustering into the 5 expected density fractions based on metabolite distributions (Supplemental Figure 4A). This indicates high replicate reproducibility, comparable to other studies (Beshir et al., 2019). Pairwise Pearson correlation coefficients verified that chorismate clustered with MEP pathway and CBC cycle intermediates DXP and S7P (*r* > 0.75), which are known to be restricted to the chloroplast (Figure 5; Supplemental Table 3). Subcellular distribution analysis based on linear regression (Hernandez et al., 2023) suggested that >70% of chorismate was associated with the chloroplast. Conversely, NAF analysis of isochorismate suggested that it primarily resides outside the chloroplast, with approximately 85% correlating with the vacuolar and cytosolic compartments (Figure 5c; Sup Table NAF). It displayed high correlation with other organic acids such glyoxylate, fumarate, and malate (*r* > 0.89), while its correlation score with chorismate, DXP and S7P was < 0.5. Shikimate correlated strongly with isochorismate (*r* ∼0.97) and vacuolar acids (*r* = 0.77 – 0.91) in wild-type plants, while still demonstrating correlations above 0.5 for plastidic markers and chorismate. It was estimated that ∼67% of shikimate may reside outside of the chloroplast (Supplemental Table 3), in agreement with its low, linear rate of ^13^C incorporation. This dual localization of shikimate is consistent with previous assessments of its localization based on NAF and isotopic labeling (Szecowka et al., 2013; Bergman et al., 2022).

When the same NAF analysis was applied to the *eds5-1* mutant, the isochorismate correlation to vacuolar acids declined from >0.9 to 0.5-0.6 (Figure 6B). In contrast, its correlation with chorismate rose from 0.38 to 0.70, although its low concentration in the mutant complicated its interpretation. Chorismate in contrast consistently correlated with plastid metabolites DXP and S7P in both genotypes. NAF analysis of *eds5-1* mutant plants displayed higher variability overall (Supplemental Figure 4B,D), with HCA analysis yielding an NMI score of 0.62 (n = 3) for clustering of the 5 fractions. This may reflect greater variation in metabolic outcomes resulting from perturbation of isochorismate trafficking. However, overall, our NAF data strongly support our initial hypothesis that chorismate is almost entirely confined to the chloroplast compartment while most isochorismate resides outside it.

### Global carbon commitment

To compare total carbon commitment into isochorismate and chorismate, we applied the external calibration curve constructed for chorismate to isochorismate based on the similarity in mass and fragmentation pattern. Comparison of total carbon commitment into chorismate and isochorismate indicated that approximately one third of the chorismate pool is diverted to isochorismate (Table 1), most of which appears exported to the cytosol based on its labeling pattern and matching NAF results. These results are consistent with the interpretation that most cytosolic isochorismate represents a pre-existing, slowly exchanging reservoir rather than a rapidly cycling intermediate. To assess whether such accumulation was unique to Arabidopsis, we compared isochorismate and chorismate levels in species known or suspected to synthesize SA via alternative routes, such as the benzoyl-CoA–mediated pathway (Liu et al., 2025; Ma et al., 2025; Wang et al., 2025; Zhu et al., 2025). In *Populus trichocarpa* and *Nicotiana benthamiana* leaves, isochorismate was detected only in trace amounts (Figure 4D and 4E). These interspecific differences support the interpretation that Arabidopsis possesses an unexpectedly large extra-plastidic isochorismate pool that potentially supplies SA production during pathogen challenge through the ICS pathway as well as other specialized routes of cytosolic aromatic metabolism (Scholten et al., 2024; Wu et al., 2024).

## Discussion

### Isotopic labeling in intact plants highlights branching and compartmentation in shikimate pathway metabolism

The present work resolves several analytical and conceptual challenges surrounding the quantification of shikimate pathway flux and transport. As the product of the shikimate pathway, chorismate is the last common intermediate leading to diverse aromatic natural products in plants (El-Azaz and Maeda, 2025). We implemented a multiplexed ^13^C-based LC–MS/MS framework to reconstruct its isotopologues unambiguously for fractional labeling calculations. Leveraging the characteristic fragmentations of chorismate, isochorismate, and shikimate, we were able to distinguish kinetic signatures of plastidial and extraplastidial carbon flow through this central aromatic pathway.

Isotopic labeling strategies have previously been applied to studying the shikimate pathway and AAA biosynthesis in plants. El-Azaz et al. recently reported time-course labeling of intact plants to highlight differential regulation of the shikimate pathway between monocots and dicots (El-Azaz et al., 2023). Using multi-hour time-course labeling experiments, the authors demonstrated that allosteric regulation of DAHP synthase evolved in grasses to accommodate the inclusion of Tyr in addition to Phe for lignin biosynthesis in that clade. Tyr:Phe ratios were further explored with labeled AAAs in wheat, which found that upon fungal elicitation, Tyr was preferentially incorporated into downstream defense metabolites over Phe (Rypar et al., 2025). The novelty of our approach rests in the measurement of the rate of appearance of ^13^C label in any metabolite pool relative to the global ^13^C intake, providing a benchmark for comparing absolute carbon commitment by metabolite that is independent of its rate of consumption. The comparison of time-resolved, targeted flux measurements paired to EA-IRMS data further enables the identification of inactive metabolite stores and infers transport processes. Thus, these experiments point to not only the fraction of total assimilated carbon entering the shikimate pathway but also the proportion that completes the sequence to chorismate.

Our calculations indicate that in Arabidopsis leaf, about one third of shikimate carbon departs the canonical sequence to chorismate and presumably accumulates outside the chloroplast. In the cytosol, shikimate participates in monolignol biosynthesis by forming transient esters with *p*-coumarate during caffeoyl-CoA formation needed for guaiacyl and syringyl lignin formation (Schoch et al., 2001; Hoffmann et al., 2003; Hoffmann et al., 2004; Saleme et al., 2017), but is evidently not turned over further, resulting in slow exchange of isotopic label. This pre-existing, unlabeled shikimate ester pool, recycled during monolignol formation with minimal net flux (Vanholme et al., 2013), may account for the low, near linear rate of ^13^C incorporation observed in whole tissue homogenates, which combine this cytosolic pool with concurrent *de novo* shikimate synthesis in the chloroplast during ^13^CO_2_ labeling. The mechanism of shikimate export from the chloroplast is currently unknown.

In contrast, chorismate appears exclusively localized to the chloroplast compartment based on its rapid, hyperbolic labeling pattern (half-life of ∼5 min) and high labeling plateau (∼70% enrichment in < 10 min). This labeling pattern parallels that of CBB cycle intermediates observed during its early elucidation, which reached only 70-80% ^14^C enrichment when fed 100% ^14^CO_2_ (Sharkey, 2019). This isotopic dilution was later shown to be due to unlabeled stores of pentose phosphates undergoing import during photosynthesis (Xu et al., 2022). The rapid formation and conversion of chorismate implies both formation and utilization close to the site of primary carbon fixation, which is inconsistent with the notion of a chorismate pool outside the chloroplast.

While hyperbolic labeling and rapid turnover are characteristic of metabolites proximal to carbon fixation (Szecowka et al., 2013), these features alone are not diagnostic of chloroplast localization. For example, PEP labels rapidly despite being produced primarily in the cytosol from triose phosphate. Its fast labeling kinetics likely reflect reimport via PPT1 in mesophyll tissue (Streatfield et al., 1999), rapid consumption by DAHPS, and low steady-state abundance (Evans et al., 2024). However, the low recovery of PEP observed here suggests its pool size may be larger than originally estimated. Conversely, chloroplast-localized metabolites with high abundance and slow turnover, such as chlorophyll *a/b* and carotenoids may require hours to days for detectable labeling (Beisel et al., 2010; Banh et al., 2022), resulting in approximately linear label incorporation during short-term assays. Thus, labeling kinetics alone do not resolve compartmentation (chloroplast vs cytosol or other extra-plastidic compartment). In this context, the combined evidence from NAF analysis, ^13^C time-course labeling, and genetic evidence obtained with the *eds5-1* mutant provide strong support for predominantly chloroplastic localization of chorismate in a brassicaceous species such as Arabidopsis. This does not exclude that chorismate may play a functional role in the cytosol of plant species engaged in specialized metabolism through the shikimate pathway, such as benzenoid volatile production in petunia petals (Qian et al., 2019).

While a parallel, cytosolic route to chorismate has been proposed (Lynch, 2022), direct evidence in a model plant system remains lacking. In Arabidopsis, most shikimate pathway enzymes, including *enol*pyruvylshikimate phosphate synthase (AT2G45300 and At1g48860) and chorismate synthase (AT1G48850), are encoded by genes predicted to produce plastid-targeted proteins with no clear cytosolic paralogs. Although plastid-targeting does not preclude cytosolic localization through alternative transcriptional and translational start sites (Willems et al., 2021), such mechanisms, while well-established in terpenoid metabolism (Cunillera et al., 1997; Phillips et al., 2008; Ruiz-Sola et al., 2016), have not yet been demonstrated for shikimate pathway enzymes. Indeed, Willems et al. detected alternative translation products for terpenoid enzymes but not those of the shikimate pathway (Willems et al., 2021).

Together, our data support predominantly chloroplast localization of chorismate in mesophyll tissue but dual compartmentation for shikimate, which partitions between plastidial and cytosolic pools with distinct turnover kinetics. However, we cannot exclude that chorismate may play a role outside the plastid in heterotrophic tissues. For instance, various chorismate utilizing enzymes, some of which are predicted to localize to the cytosol, produce a series of related non-aromatic, isomeric compounds in the roots of Arabidopsis (Peng et al., 2026).

### Fluxes into aromatic and terpenoid precursor pathways are comparable in photosynthetic tissue

The integrated labeling and flux analyses demonstrate that, in mature Arabidopsis leaves, the shikimate pathway operates at unexpectedly low flux compared to the plastidic MEP pathway, consuming only ∼0.05% of newly assimilated carbon in photosynthetic tissue. This challenges the notion that 30% of photosynthetically fixed carbon passes through Phe in plant systems generally (Haslam, 1993; Razal et al., 1996; Boerjan et al., 2003; Qian et al., 2019) and suggests that carbon allocation into phenolic metabolic pathways depends on tissue type. While woody species, (El-Azaz and Maeda, 2025), lignin-rich vascular tissue, or tissue specializing in phenylpropanoid volatile production (Boatright et al., 2004; Maeda et al., 2010; Lynch et al., 2017; Qian et al., 2019) may indeed direct a substantial portion of fixed carbon to Phe biosynthesis, our results suggest that the fraction of total assimilated carbon dedicated to phenolic precursors in photosynthetic mesophyll tissue is well below this figure. This may reflect the diminished requirement for shikimate pathway products in mesophyll tissue, compared to the elevated need for Phe for lignin precursors in vascular tissue and stems (Ehlting et al., 2005), both of which consume large quantities of PEP for lignification (Evans et al., 2025).

The large discrepancy in shikimate pathway flux between lignin-rich tissue and mesophyll tissue may depend on both biochemical factors and substrate supply. For instance, in *Petunia x hybrida* petals, downregulation of arogenate dehydratase, leading to Phe, unexpectedly resulted in an overall decrease in flux to shikimate (Maeda et al., 2010). Certain AAA isoforms are allosteric inhibitors of DAHPS, the first committed step of the pathway (Yokoyama et al., 2021). Elevated Tyr levels cause the cytosolic retention of Arabidopsis DAHP2 isoform through interaction with a 14-3-3 protein (Kanaris et al., 2022), possibly providing another means to reduce shikimate pathway flux in tissues where its requirement is reduced, although the details remain unclear.

Recent progress toward understanding the regulation of the shikimate pathway may help explain the bifurcation of shikimate between compartments. Arabidopsis mutants defective in the TyrA gene encoding arogenate dehydrogenase display a stunted growth and reticulated leaf phenotype (de Oliveira et al., 2019). A genetic screening for TyrA suppressor mutants identified multiple lines mapping to DAHPS (Yokoyama et al., 2022). These *suppressor of tyra* (*sota*) mutants carry point mutations that abolish feedback inhibition by AAA, resulting in increased CO_2_ assimilation, elevated AAA levels, and higher flux through the shikimate pathway, as reflected by increased ^13^C labeling rates and expansion of the shikimate pool (Yokoyama et al., 2022). In terms of metabolic control theory (Fell, 1997), DAHPS isoforms normally exhibit negative elasticities towards pathway end products, analogous to DXS in the MEP pathway, which is inhibited by IDP and DMADP (Banerjee et al., 2013; Wright et al., 2014; Mitra et al., 2021). Negative product elasticity at the first committed step is a common network feature that dampens fluctuations in metabolite pools while permitting modulation of pathway flux. Loss of this feedback in *sota* mutants removes a stabilizing constraint, increasing pathway flux and reducing concentration buffering, thereby allowing downstream metabolites such as shikimate to accumulate. These observations are consistent with a substantial, metabolically responsive chloroplast-localized pool of shikimate that can expand when flux constraints at DAHPS are relieved, even if a significant fraction of total cellular shikimate resides outside the chloroplast.

Beyond regulation at the biochemical level, restriction of substrate supply may also directly impact flux. Reimport of PEP into chloroplasts is governed by two PEP:phosphate transporters (PPT1 and PPT2). PPT1 is broadly expressed in vascular tissues and roots, resulting in a distinct reticulated phenotype when mutated (Streatfield et al., 1999; Voll et al., 2003), whereas PPT2 is primarily expressed in mesophyll tissue but does not complement the PPT1 mutant phenotype (Knappe et al., 2003), also known as the *chlorophyll a/b binding protein underexpressed* (*cue1*) mutant. The *cue1* phenotype has been speculated to derive from more than a simple lack of AAA (Voll et al., 2003), and the cell-specific requirement for high shikimate flux in vascular tissue for lignification, where PPT1 is highly expressed, is consistent with the notion that disruptions in phenylpropanoid availability for lignin biosynthesis, not AAA shortage for protein synthesis, is the source of the reticulated phenotype of *cue1* (Staehr et al., 2014).

Although the total carbon commitment to shikimate pathway is lower than expected, the total carbon investment into the MEP pathway is similar. Although we observed <1% of assimilated carbon entering the MEP pathway, our short term labeling approach captures flux only through rapidly turning-over intermediates and does not resolve incorporation rates into different downstream end products, including photosynthetic pigments, prenylated lipids, and signaling metabolites (Ramel et al., 2012; Nisar et al., 2015). Isotopic labeling studies indicate that these compounds turn over on a timescale of hours (Beisel et al., 2010) to days (Banh et al., 2022), well beyond the timeframe of the precursor pathways described here. Future metabolic studies to extend the carbon commitment framework to functionalized terpenoid end products must overcome several physical limitations. Among them, their slow biosynthetic and turnover rates; the presence of large, pre-existing pools whose low natural abundance dilute early isotopic enrichment; and their structural complexity, particularly for chlorophyll, which obtains less than half its carbon from the MEP pathway. Consequently, quantitative studies to track flux into photosynthetic pigments using a similar framework will require both specialized labeling strategies and development of appropriate analytical methods.

### The cytosolic isochorismate pool may serve as a reservoir for defense-related carbon resources

Shikimate and isochorismate, but not chorismate, highlight export of pre-aromatic intermediates in mesophyll cells. Extra-plastidic isochorismate constitutes a large, metabolically inert reservoir (approximately 50-fold higher in magnitude than chorismate) with slow isotopic labeling kinetics, suggesting limited turnover under basal conditions. This pool may represent a previously unrecognized carbon reserve linking the shikimate pathway to inducible defense metabolism.

High cytosolic isochorismate levels imply that pre-existing isochorismate could potentially provide an immediate substrate for SA biosynthesis during pathogen attack or abiotic stress. If accessible to cytosolic SA-forming enzymes such as PBS3 or other yet-unidentified N- glutamyl-isochorismate lyases, this pool could enable rapid SA accumulation within minutes, prior to de novo induction of biosynthetic enzymes. Moreover, SA accumulation is not PBS3/EPS1 dependent in Arabidopsis (Lefevere et al., 2020), highlighting previous observations that isochorismate-independent sources of SA are also important (Huang et al., 2010). However, ICS induction remains important to sustain and amplify SA production (Wildermuth et al., 2001). Transcriptional upregulation of ICS and EDS5 (Nawrath et al., 2002) would provide a secondary, sustained phase of SA biosynthesis over a timescale of hours to days (Yang et al., 2015; Zheng et al., 2015; Vlot et al., 2021). Mechanistically and evolutionarily, a two-phase response (fast pool- driven production followed by transcriptionally reinforced synthesis) holds obvious advantages for the plant, and ICS induction alone does not exclude formation from pre-existing reserves of isochorismate in the cytosol. However, the question of whether SA biosynthesis from preformed precursors occurs independently of *de novo* production in Arabidopsis has not yet been addressed with the appropriate methodology (e.g., isotopic labeling with pathogen induction at high temporal resolution). In a related species, rapid SA accumulation within minutes was observed in *Brassica rapa* in response to abiotic stress (Ang et al., 2024), and the kinetics suggest that formation from pre-existing substrate/enzyme pools is at least feasible.

In addition to its role as precursor to SA, isochorismate serves as a substrate for cytosolic aromatic metabolism in Arabidopsis, including the formation of O-substituted benzoates involved in defense (Scholten et al., 2024), and alternative AAA pathways (Wu et al., 2024). In the latter, the cytosolic aromatic aminotransferase, REVERSAL OF SAV3 PHENOTYPE 1 (VAS1), channels isochorismate, following its converson to isoprephenate, into carboxylated AAA intermediates (e.g., 3-carboxy-4-hydroxyphenylpyruvate or 3-carboxylphenylpyruvate), which are further transaminated to 3-carboxy-Tyr and 3-carboxy-Phe (Wu et al., 2024). These pathways rely on EDS5-dependent export of isochorismate to the cytosol but diverge downstream of isoprephenate, illustrating their role as precursors in cytosolic metabolism of aromatic compounds. Together, these findings support an expanded function for cytosolic isochorismate in both defense and modulation of AAA biosynthesis. Comparative analyses across species further indicate that this feature may be specific to Arabidopsis and closely related taxa employing the ICS-dependent route to SA (Hong et al., 2025).

Together, these findings provide a revised view of the shikimate–chorismate– isochorismate axis as a dynamically regulated interface between plastidial and cytosolic branches of aromatic metabolism and defense signaling. Their distinct kinetic signatures and carbon allocation patterns furthermore highlight the previously unrecognized importance of isochorismate outside the chloroplast. Tandem mass spectrometry combined with structurally diagnostic CID fragmentation enabled quantitative tracking of chorismate isotopologues and their phenolic precursors and derivatives, providing a robust framework for studying aromatic metabolism. Isotopic flux analyses in Arabidopsis plants exposed to ^13^CO_2_ revealed that chorismate synthesis and turnover are likely confined to the chloroplast, whereas shikimate and isochorismate are exported and maintain substantial extra-chloroplastic pools. Notably, cytosolic isochorismate accumulates to high levels and turns over slowly, suggesting a role as a latent carbon reservoir linking primary metabolism with inducible defense pathways. Follow up studies will focus on these novel roles of aromatic precursors at the interface of primary and secondary metabolism.

---

**Supplemental Table 1.**
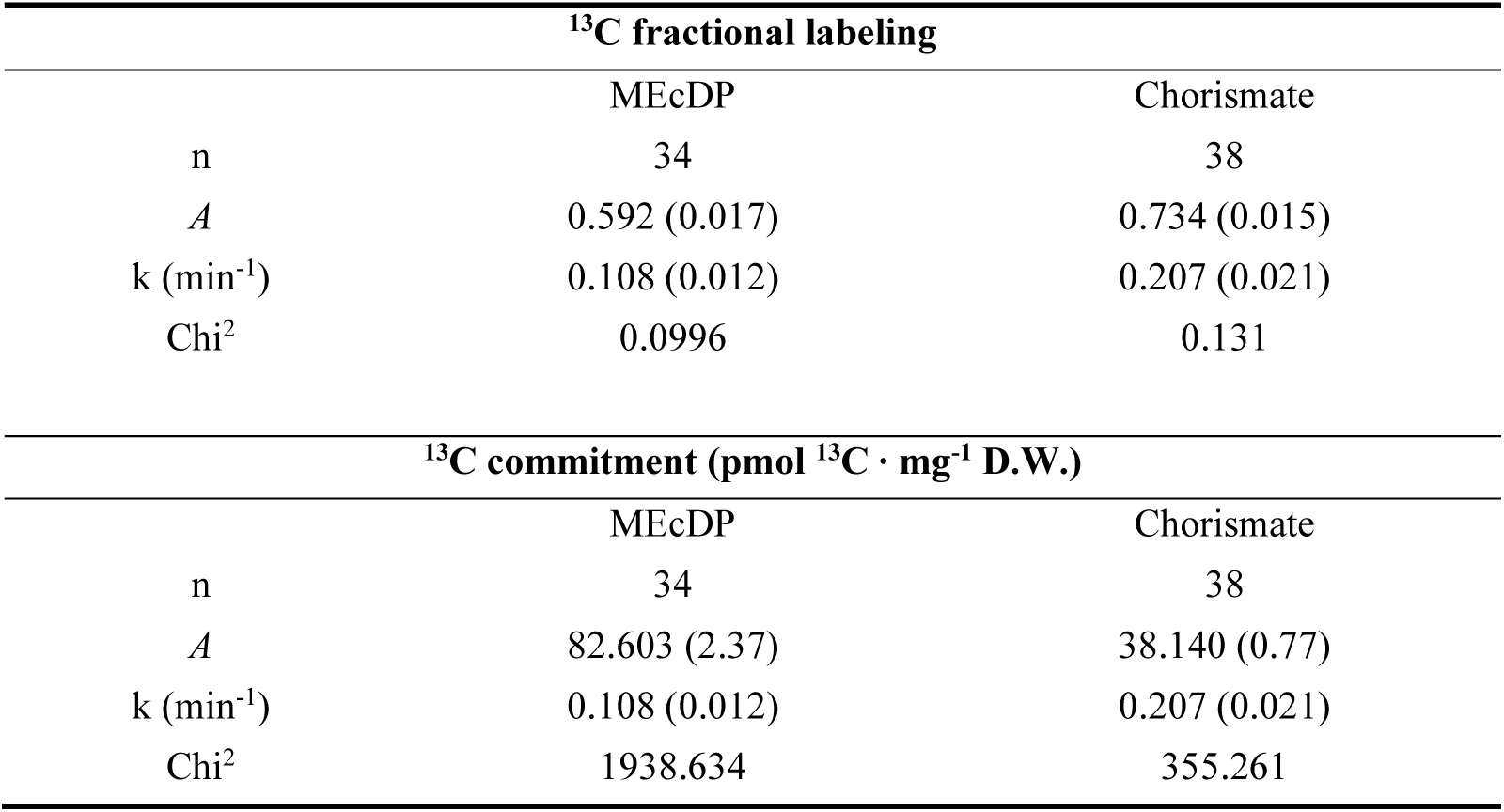
Kinetic parameters of ^13^C fractional labeling and carbon commitment for each metabolic pool. Data were fit to an exponential rise to maximum equation *A* × (1 - *e ^-k^ ^t^*), where A represents the labeling plateau, t is the labeling time, and k is the kinetic rate constant. Chi² values represent the unweighted sum of squared residuals between the observed and fitted data. Standard errors are given in parentheses. Shikimate, isochorismate, and sucrose labeling data could not be fit to the exponential rise to maximum model as ^13^C incorporation into these pools was approximately linear in plants labeled from 0 – 154.8 mins.

| $^{13}\text{C}$ fractional labeling | | |
| --- | --- | --- |
|  | MEcDP | Chorismate |
| n | 34 | 38 |
| A | 0.592 (0.017) | 0.734 (0.015) |
| k (min<sup>-1</sup>) | 0.108 (0.012) | 0.207 (0.021) |
| Chi<sup>2</sup> | 0.0996 | 0.131 |
| $^{13}\text{C}$ commitment (pmol $^{13}\text{C} \cdot \text{mg}^{-1}$ D.W.) | | |
|  | MEcDP | Chorismate |
| n | 34 | 38 |
| A | 82.603 (2.37) | 38.140 (0.77) |
| k (min<sup>-1</sup>) | 0.108 (0.012) | 0.207 (0.021) |
| Chi<sup>2</sup> | 1938.634 | 355.261 |

**Supplemental Table 2.**
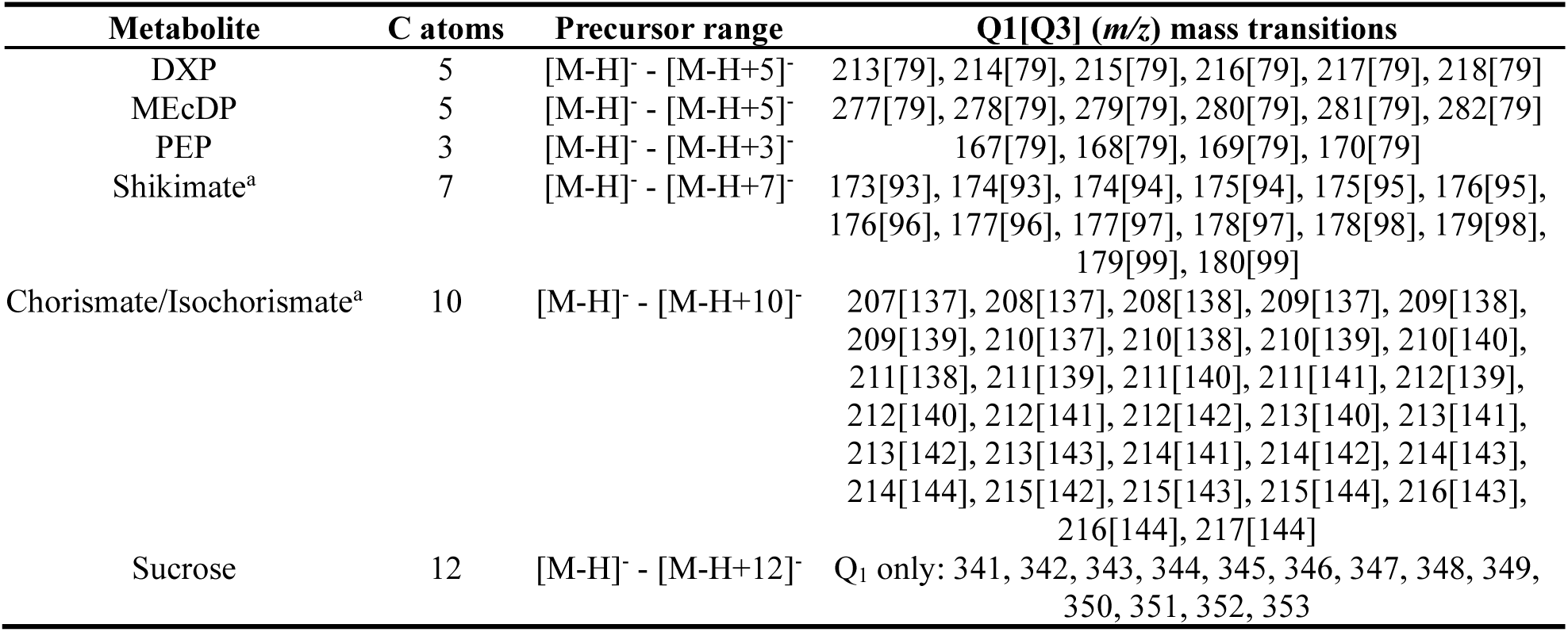
Mass isotopologues used in the calculation of fractional labeling of each metabolite pool. Shikimate, chorismate, PEP and MEP pathway intermediates were analyzed by liquid chromatography – tandem mass spectrometry (LCMS/MS) using a Sciex 4500 Qtrap operating in multiple reaction monitoring (MRM) in negative mode. Q1, *m/z* of precursor ion; Q3, *m/z* of product ion. Sucrose was analyzed using a selected ion monitoring method in Q1.

**Supplemental Table 3.** Average subcellular distribution of selected metabolites in *Arabidopsis*. Subcellular distributions were predicted using the NAFalyzer R shiny app (https://github.com/cellbiomaths/NAFalyzer), based on analysis of select metabolites in fractions obtained through non-aqueous fractionation. Abundance of DXP and UDP-glucose were used as markers for the chloroplast and cytosol respectively, while the average abundance of malate, fumarate and citrate was used as a marker for the vacuole. Data represent the mean ± SE (n=3). Abbreviations: S7P; sedoheptulose-7-phosphate.

| Metabolite | Chloroplast (%) | Cytosol (%) | Vacuole (%) |
| --- | --- | --- | --- |
| Shikimate | 23.05 ± 2.74 | 41.30 ± 5.71 | 35.67 ± 6.40 |
| Chorismate | 70.03 ± 3.40 | 28.92 ± 2.83 | 1.05 ± 0.56 |
| Glyoxylate | 26.42 ± 3.81 | 24.30 ± 13.18 | 49.28 ± 9.30 |
| Isochorismate | 14.86 ± 0.49 | 35.57 ± 3.71 | 49.57 ± 3.19 |
| S7P | 87.57 ± 6.47 | 11.92 ± 6.44 | 0.51 ± 0.16 |

**Supplemental Table 4.**
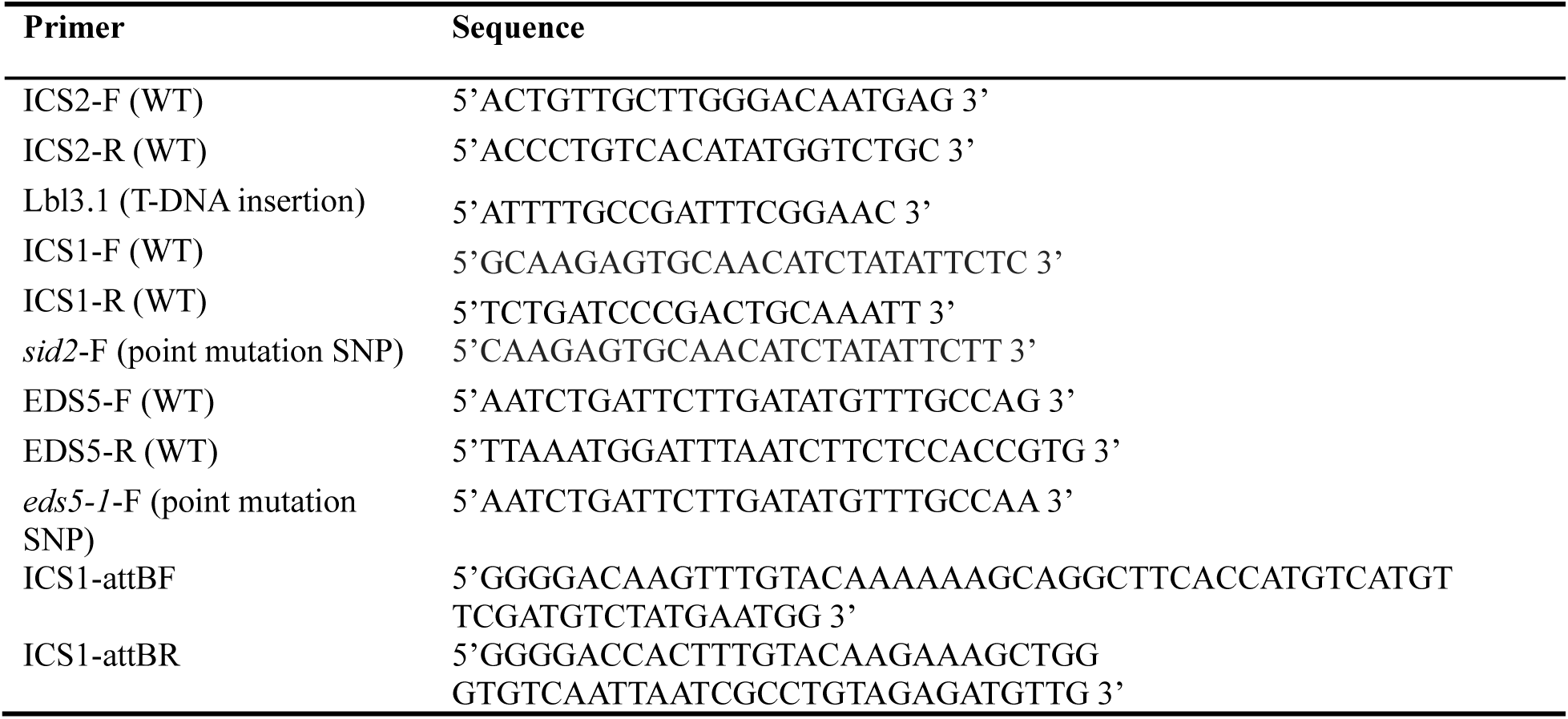
Primers used in this study. Genotyping primers for T-DNA insertion lines (*ics2)* were screened with WT primers in combination with a primer annealing to the T-DNA sequence to determine homozygosity. *Sid2* and *eds5-1* mutants contained point mutations at the 3’ position and were screened with WT and mutant primers that differed by the 3’ base position. Cloning was performed using the Gateway^TM^ cloning system with primers containing attB adaptors.

**Supplemental Table 5.** MS parameters used for metabolite quantification by LC-MS/MS multiple reaction monitoring (MRM)

| Metabolite | Precursor | Product | ESI<br>polarity | DP<sup>a</sup> | EP<sup>a</sup> | CE<sup>a</sup> | CXP<sup>a</sup> |
| --- | --- | --- | --- | --- | --- | --- | --- |
|  | (m/z) | (m/z) |  | (V) | (V) | (V) | (V) |
| DXP | 213 | 79 | (-) | -30 | -10 | -40 | -8 |
| MEcDP | 277 | 79 | (-) | -30 | -10 | -65 | -11 |
| PEP | 167 | 79 | (-) | -5 | -10 | -14 | -5 |
| Shikimate | 173 | 93 | (-) | -51 | -10 | -19 | -10 |
| Chorismate/Isochorismate | 207 | 137 | (-) | -53 | -4 | -16 | -4 |
| S7P | 289 | 96.9 | (-) | -38 | -10 | -22 | -7 |
| UDP-gluc | 565 | 323 | (-) | -45 | -8 | -36 | -7 |
| Malate | 133 | 115 | (-) | -30 | -10 | -18 | -4 |
| Fumarate | 115 | 71 | (-) | -32 | -8 | -13 | -2 |
| Succinate | 117 | 73 | (-) | -41 | -10 | -18.5 | -2 |
| Glyoxylate | 73 | 45 | (-) | -50 | -9 | -11.2 | -5 |
| Citrate | 191 | 111 | (-) | -30 | -10 | -18 | -4 |
| DGP<sup>b</sup> | 243 | 79 | (-) | -35 | -10 | -62 | -7 |

<sup>a</sup> ESI, electrospray ionization; DP, declustering potential; EP, entrance potential; CE, collision energy; CXP, collision cell exit potential; V, volts
<sup>b</sup> 2-deoxyglucose-6-phosphate (DGP), used as internal standard for phosphorylated metabolite quantification

## Acknowledgements

This study was funded by a Discovery grant from the Natural Sciences and Engineering and Research Council (NSERC) of Canada (RGPIN-2024-05076) and a John Evans Leadership Fund grant from the Canadian Foundation for Innovation (36131). The authors acknowledge a generous NSERC CGSM graduate scholarship supporting AEF and SAF and a CGSD graduate scholarship supporting MEB.

## Statement of data availability

## Author contributions

S.A.F. and A.E.F. performed main experiments. A.H., C.D.M., and S.E.E. performed minor experiments. M.A.P. wrote the manuscript and directed the research.

---

**Supplemental Figure 1.**
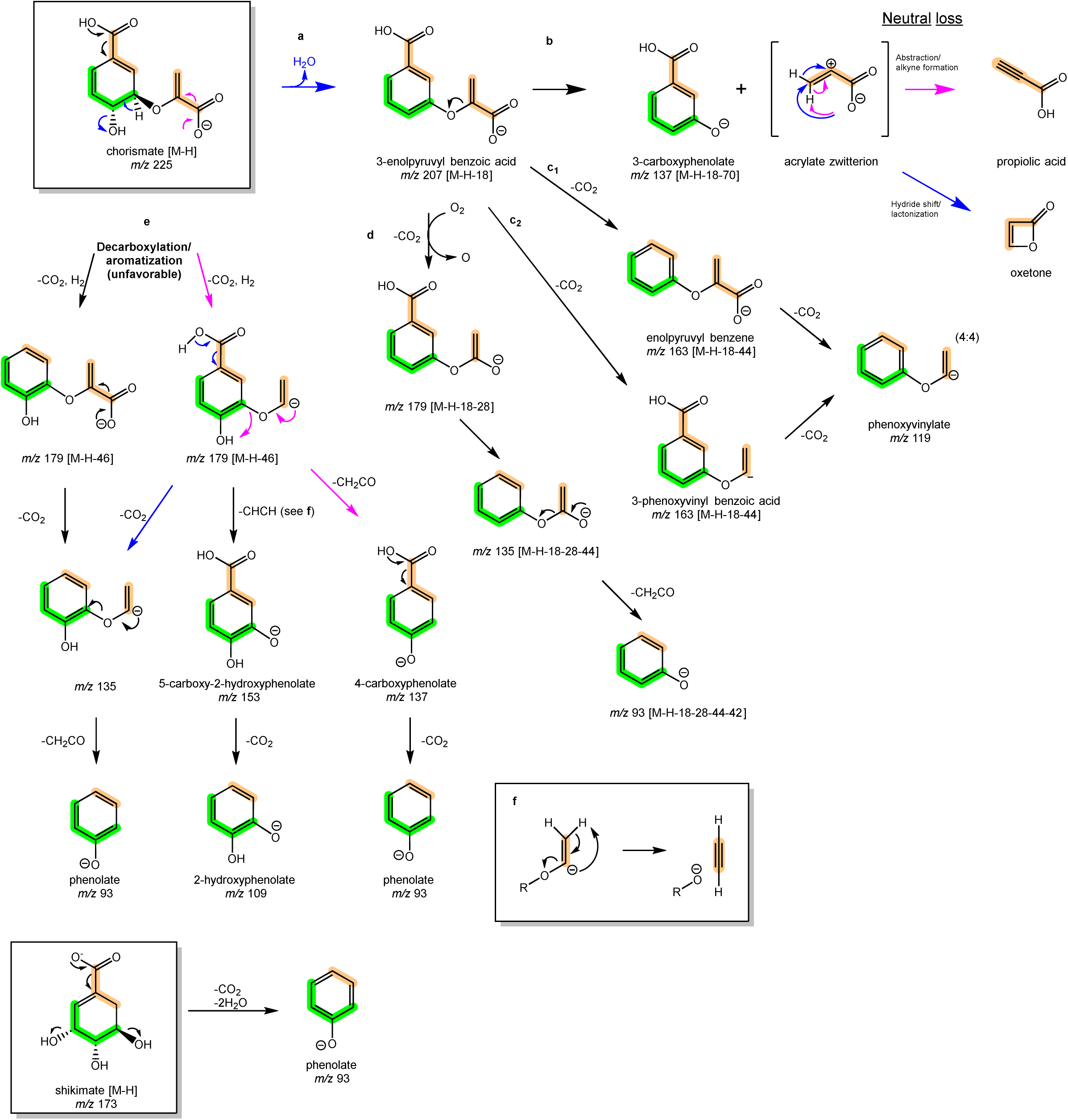
Proposed fragmentation of chorismate and shikimate during collision induced dissociation (CID) and MS^3^ analysis using direct infusion of purified standards. Carbons derived from D-erythrose-4-phosphate (E4P) are highlighted in green while those obtained from phosphoenolpyruvate (PEP) are shown in orange. The chorismate quasimolecular [M-H]^-^ ion (*m/z* 225) is unstable and undergoes two forms of in source fragmentation. In the first, dehydration forms 3-enolpyruvyl benzoate (**a**). MS^2^ and MS^3^ scans suggest the 207 dehydration product fragments by 3 distinct paths: Loss of an acrylate zwitterion ion, which rearranges to propiolic acid (**b**), loss of two CO_2_ molecules in either order, leading to a phenoxyvinylate ion (**c_1_** and **c_2_**), and formation of an O_2_ adduct following an initial CO_2_ loss, followed by decay of the resulting peroxide (**d**) for a net loss of 28 Da *(m/z* 207 → 179). MS^2^ and MS^3^ scans also suggest that aromatization can take place through decarboxylation and loss of 2H•, leading to a pair of substituted phenol rings at low efficiency (**e**), which may further undergo loss of CO_2_, ketene, or acetylene (**f**) to yield 5-carboxy-2- hydroxyphenolate (*m/z* 153), 2-hydroxyphenolate (*m/z* 109), and phenolate *(m/z* 93).

**Supplemental Figure 2.**
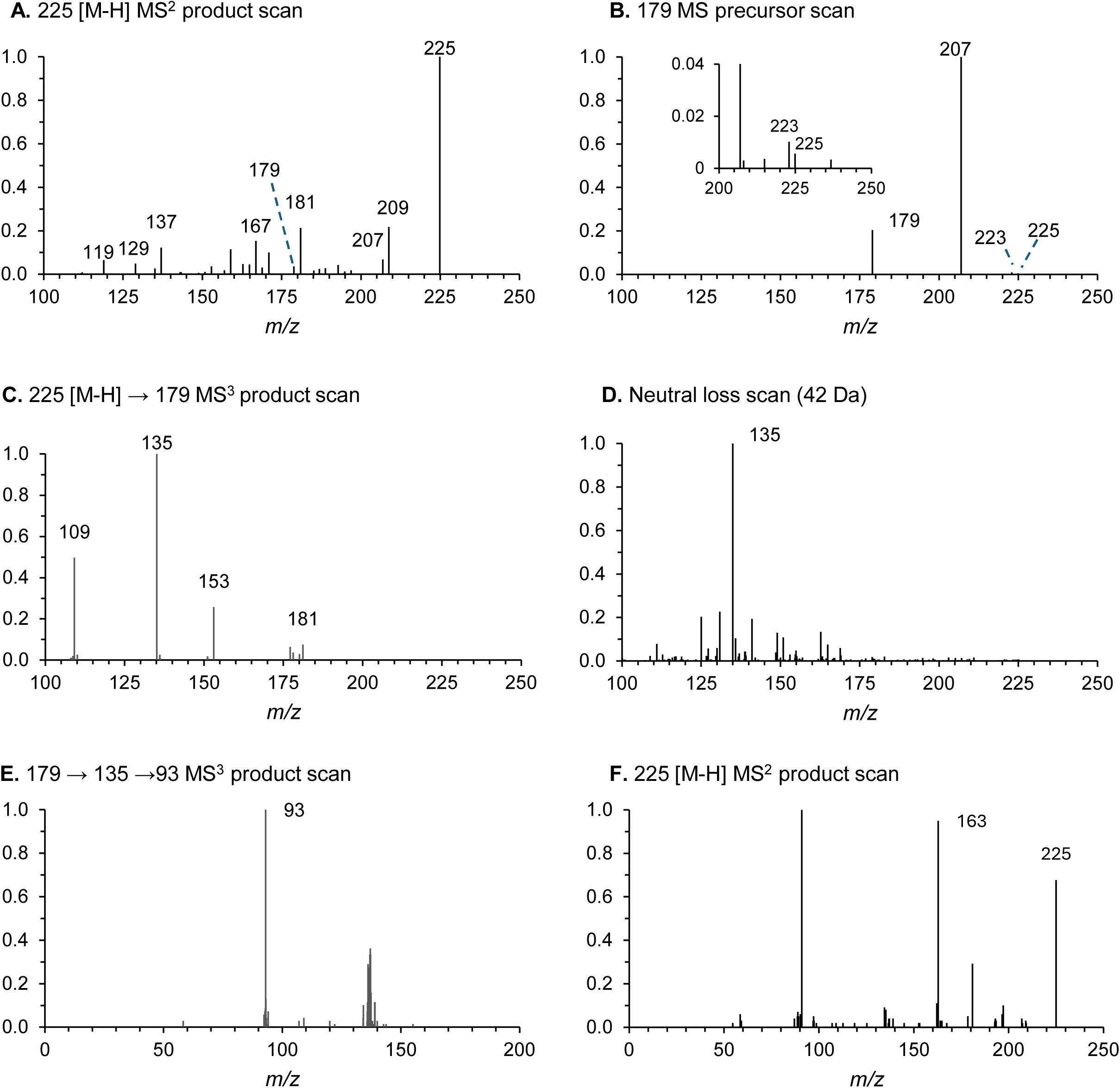
Additional MS^2^ and MS^3^ spectra of chorismate. **A**, MS2 product spectrum of *m/z* 225 [M-]. **B**, Precursor spectrum of the *m/z* 179 product ion. C, MS3 product spectrum of the m/z 225 →179 transition. **C**, MS3 spectrum of the m/z 225 [M-H] → 179 [M-H-CO2-2H]^-^ series. D, Neutral loss scan for 42 Da, confirming that *m/z* 93 product ion (*m/z* 179 → 135 →93) is the results of ketene loss from *m/z* 135. E, MS^3^ scan of the *m/z* 179, demonstrating its sequential loss of CO_2_ (*m/z* 179 → 135) and ketene (C=C=O) to form a phenolate ion (*m/z* 93), as shown in Supplemental Figure 2. F, MS^2^ product scan for the 225 prephenate [M-H] molecular ion.

**Supplemental Figure 3.**
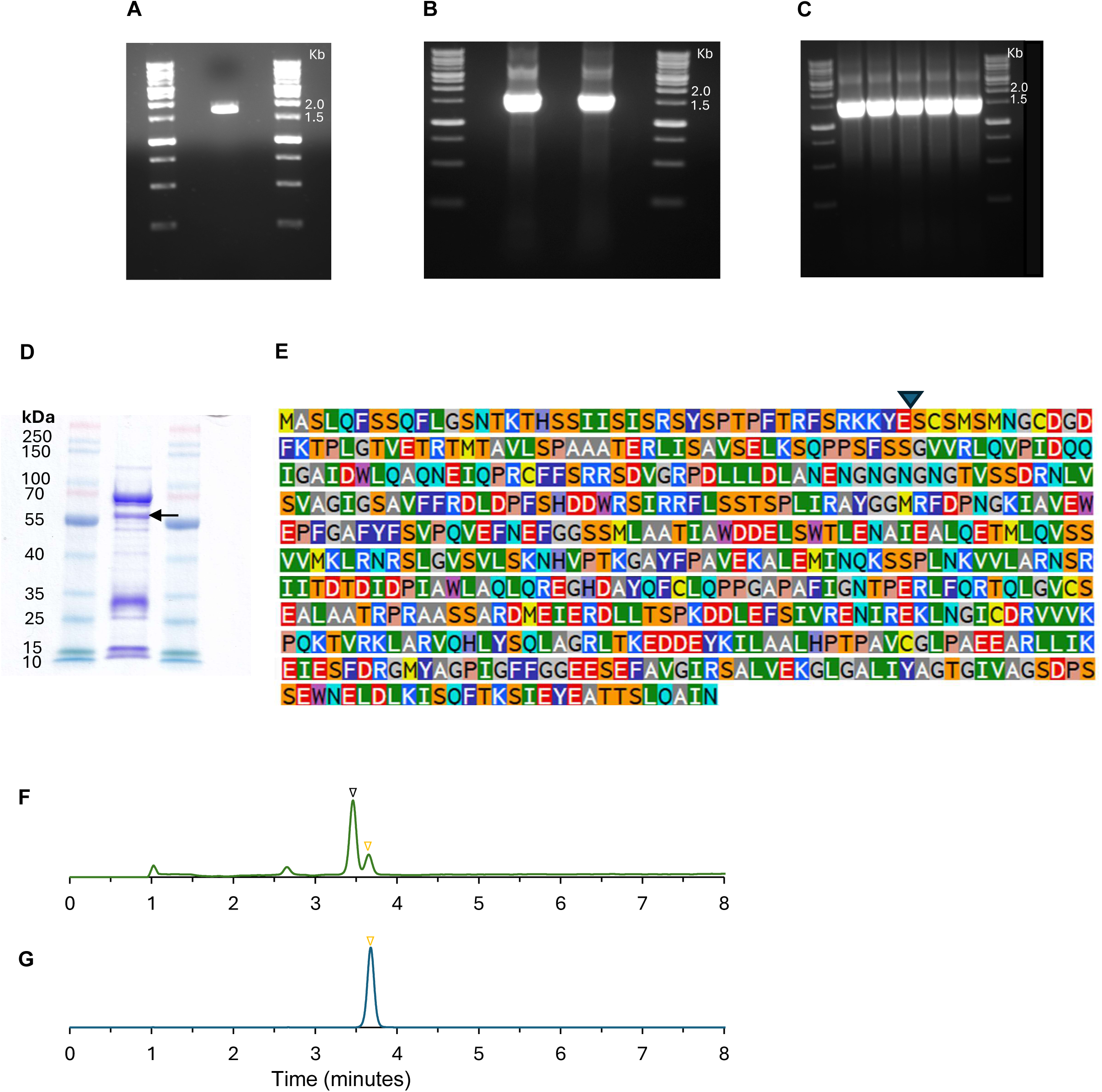
Cloning and expression of Arabidopsis isochorismate synthase 1 (ICS1). **A,** Initial PCR amplification of ICS1 with Gateway^TM^ cloning primers to generate an amplicon lacking the plastid transit peptide of ICS1 (expected size ∼ 1585 bp). **B**, Colony PCR of two colonies of *Escherichia coli* TOP10 strain transformed with the truncated ICS1 following BP Clonase cloning into pDONR207. **C**, PCR of five resistant TOP10 colonies following the LR subcloning reaction into pET28 and bacterial transformation. **D**, SDS-PAGE showing partial purification of truncated, recombinant ICS1 (indicated by arrow) (57.8 kDa). **E**, Amino acid sequence of ICS1 indicating the truncation site (blue arrow) for truncation of the chloroplast target peptide. **F**, Liquid chromatography – tandem mass spectrometry (LCMS/MS) analysis of an ICS assay using 0.8 ug of purified ICS incubated with 10 μM chorismate as substrate for 1 hour at 25°C. G. Chorismate incubated without protein (negative control) and analyzed by the same LCMS/MS method. The y axis displays the detector response corresponding to the negative mode *m/z 2*07→137 mass transition (counts per second), which represents the in-source dehydration product of the *m/*z 225 pseudomolecular ion of either chorismate or isochorismate, which share the same mass and undergo the same mass transition. Black triangle = isochorismate, yellow triangle = chorismate.

**Supplemental Figure 4.**
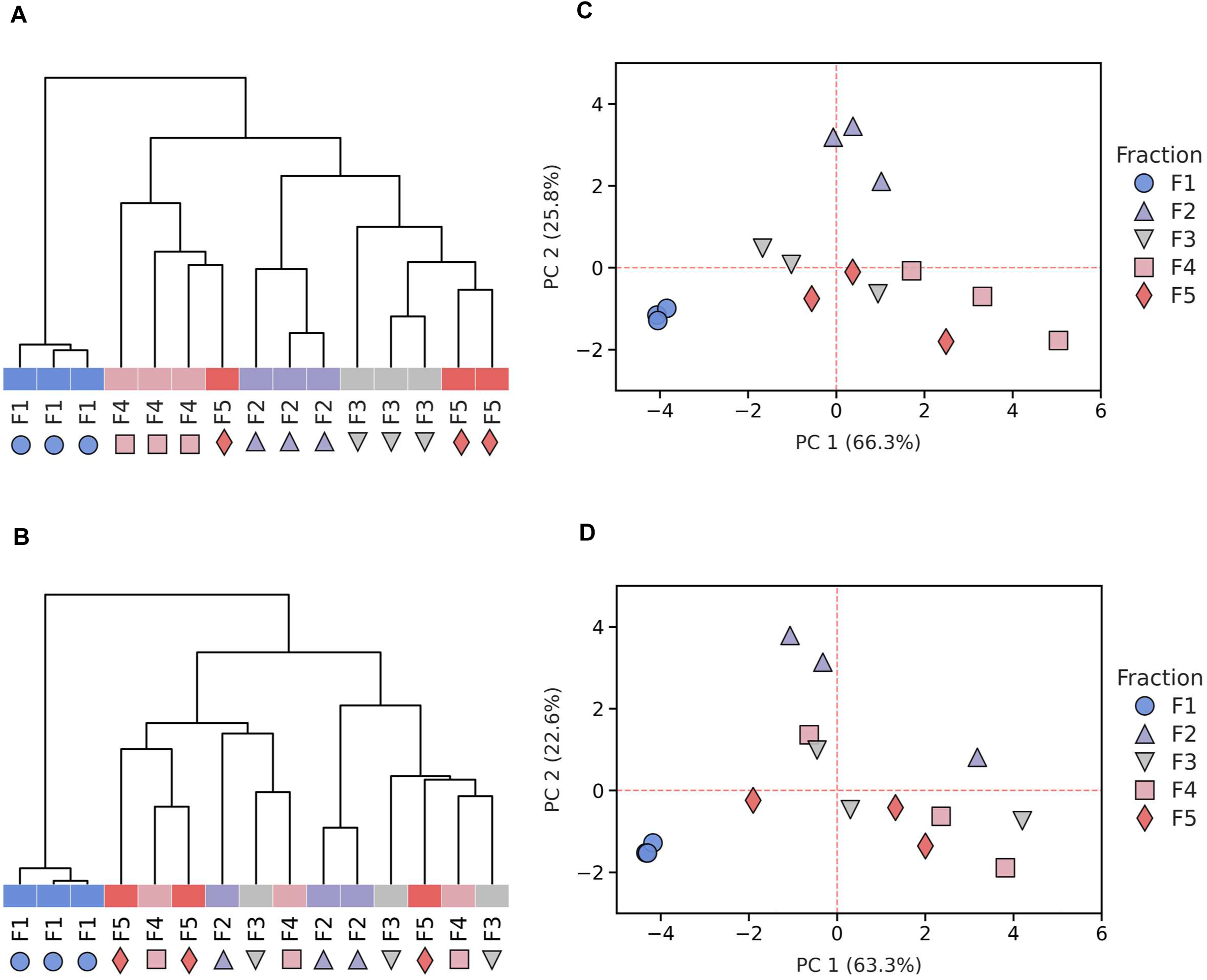
Resolution and reproducibility of non-aqueous fractionation fractions for wildtype (A, C) and the *eds5* mutant (b, d). **A, B.** Hierarchical cluster analysis (HCA) for the three biological replicates based on scaled relative abundances of individual metabolites across the five fractions obtained through non-aqueous fractionation Legend entries F2, F2, F3, F4, F5, and F6 denote fractions with densities of approximately 1.33, 1.42, 1.50, and >1.62, respectively, while F1 denotes densities <1.33 (see Methods). Heat map colors encode equivalent fractions from different replicates. The Normalized Mutual Information score of 0.81 for wild-type indicates high reproducibility across replicates (Beshir et al., 2019; See Methods). **B, D.** Principal component analysis (PCA) scores plot based on the scaled distribution of individual metabolites across fractions of different densities (n=3).

**Supplemental figure 5.**
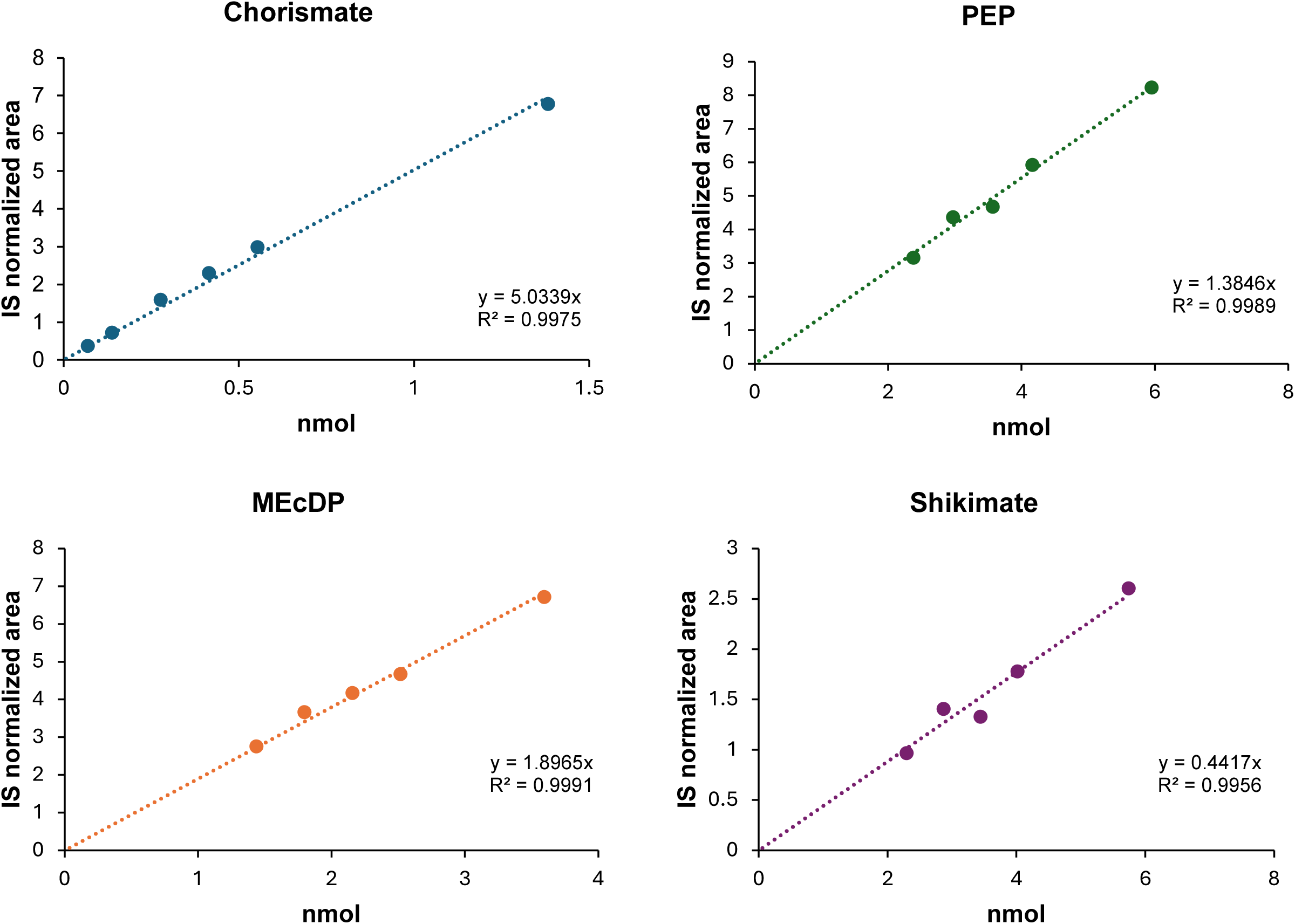
External standard calibration curves. Solvent only calibration curves prepared from spiking varying ng amounts (500-1000ng) of purified analytical standard to the initial extraction solvent, with internal standards added, and processing with the polar metabolite extraction protocol. Ratios of analyte area to the internal standard relative to ng amounts yielded the slope used to calculate pool size of chorismate and isochorismate from n=10 biological WT Arabidopsis replicates.

**Supplemental figure 6.**
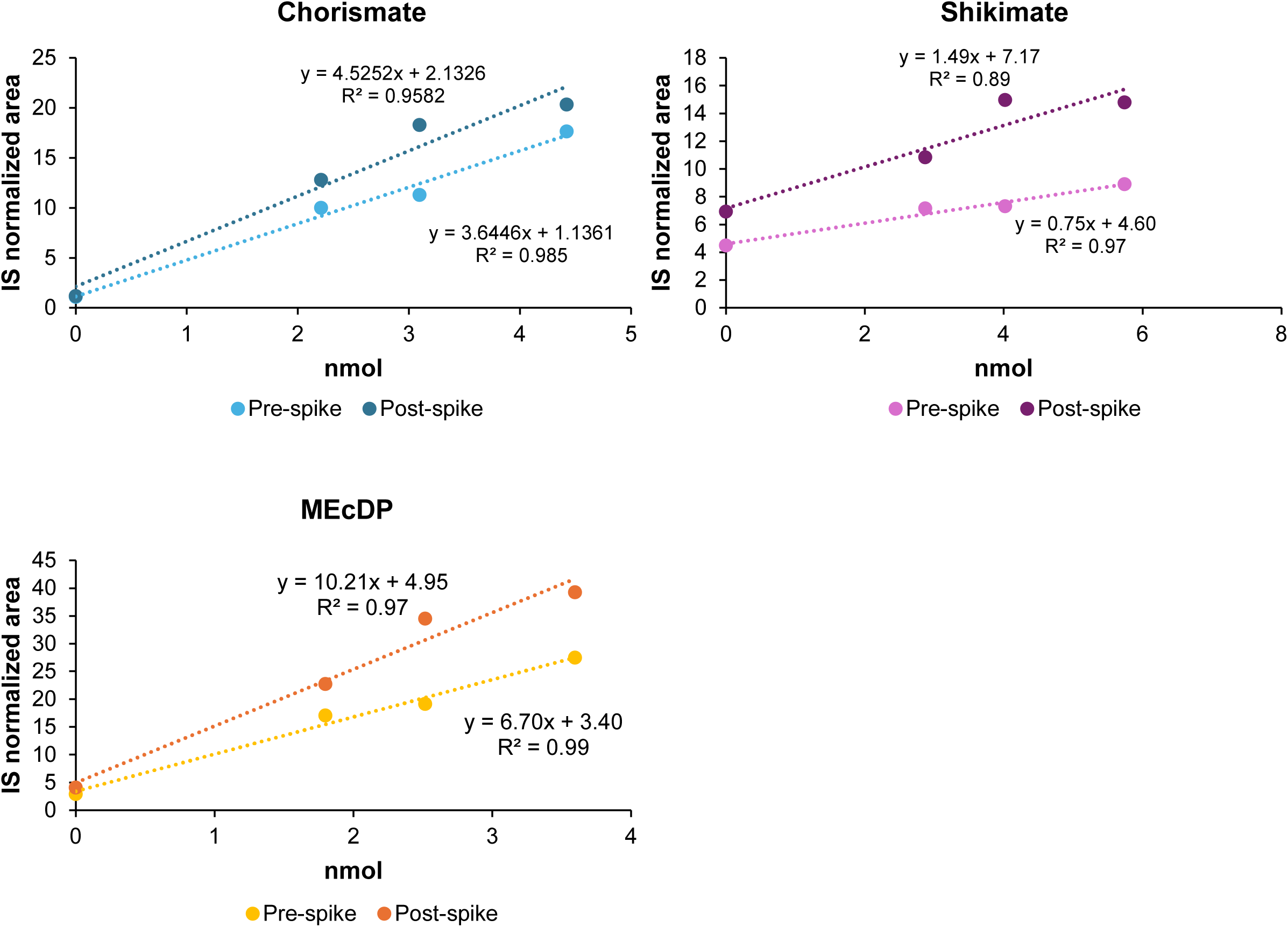
**Standard addition calibration curves**. Control polar metabolite Arabidopsis leaf tissue extracts were spiked with varying ng amounts of purified analytical standard (500-1000ng) at either the initial extraction (pre-spike) or before injection (post-spike). Differences in normalized peak areas were used to estimate recovery of metabolites due to matrix effects and protocol-specific losses.

